# Soil N_2_O emission potential falls along a denitrification phenotype gradient linked to differences in microbiome, rainfall and carbon availability

**DOI:** 10.1101/2020.07.22.211268

**Authors:** Matthew P. Highton, Lars R. Bakken, Peter Dörsch, Steve Wakelin, Cecile A. M. de Klein, Lars Molstad, Sergio E. Morales

## Abstract

Soil denitrification produces the potent greenhouse gas nitrous oxide (N_2_O) and by further reduction of N_2_O, the harmless inert gas N_2_. N_2_O emission is determined by rate and timing of the N_2_O producing and reducing steps which are sensitive to a series of proximal and distal regulators such as pH and microbial community composition. Microbial community associations to N_2_O emission potential (N_2_O/(N_2_O+N_2_)) are commonly entangled with pH leaving the true role of community composition unclear. Here, we leverage a set of soil microbiomes strongly linked to rainfall above pH to test the hypothesis that microbiome vs. N_2_O emission potential (N_2_O/(N_2_O+N_2_)) correlations will be maintained across alternative distal drivers. N_2_O emission potential (N_2_O/(N_2_O+N_2_)) and denitrification gas (NO, N_2_O, N_2_) kinetics were assessed by automated gas chromatography while community composition was assessed by 16S rRNA gene sequencing and qPCR of *nosZI* and *II* genes. Analyses revealed a sustained correlation between microbiome and N_2_O emission potential (N_2_O/(N_2_O+N_2_)) in the absence of a pH effect. Further, a continuum of gas accumulation phenotypes linked to NO accumulation and sensitive to carbon addition are identified. Separate phenotypes carried out N_2_O production and reduction steps more concurrently or sequentially and thus determined N_2_O accumulation and emission potential (N_2_O/(N_2_O+N_2_)). Concurrent N_2_O producing/reducing soils typically contained NO accumulation to a low steady state, while carbon addition manipulations which increased NO accumulation also increased sequentiality of N_2_O production/reduction and thus emission potential (N_2_O/(N_2_O+N_2_)). These features may indicate a conserved NO inhibitory mechanism across multiple effectors (rainfall, community composition, carbon availability).

**Highlights:**

- N_2_O emission potential is linked to microbiome changes associated with rainfall, but not to pH.
- Sequential vs. concurrent denitrification phenotypes differing in NO and N_2_O accumulation are identified.
- High N_2_O accumulation is associated with increased NO accumulation.
- Sequentiality of N_2_O production/reduction determines soil N_2_O emission potential.
- Sequentiality of N_2_O reduction was susceptible to manipulation via carbon addition.

## Introduction

Production and emission of nitrous oxide (N_2_O) represents a significant climate concern due to its high global warming potential (298 times that of CO_2_ over a 100 year time span on a mass to mass basis) (Myhre *et al.*, 2013) and ozone depleting activity (Ravishankara *et al.*, 2009). The most recent IPCC report ranks N_2_O as the third most significant greenhouse gas, accounting for 6.2% of global climate forcing (Intergovernmental Panel on Climate Change, 2013). Atmospheric concentrations of N_2_O have risen dramatically over the past century to a current concentration of greater than 333 ppb (Jan, 2020; 2° Institute, 2016), much of which is attributed to anthropogenic soil emissions (Davidson, 2009). Global N_2_O budgets suggest that around 45% of the emitted N_2_O is produced anthropogenically with the majority (60%) coming from agricultural sources (Syakila and Kroeze, 2011). In an agricultural setting, N_2_O production is traditionally attributed to denitrification and nitrification (Bremner, 1997) of N in animal excreta or applied fertilizers (Davidson, 2009; Syakila and Kroeze, 2011; Oenema *et al.*, 2005) but a number of other biological processes are also relevant (Baggs, 2011).

Denitrification occurs under anoxic conditions when microbial populations switch from O_2_ based respiration to reduction of nitrogenous molecules (NO_3_^-^→ NO_2_^-^ → NO → N_2_O → N_2_). In each step, reduction of the nitrogenous molecule as a terminal electron acceptor is catalyzed by an independent reductase enzyme (nitrate reductase-Nar or Nap, nitrite reductase-Nir, nitric oxide reductase-Nor, and nitrous oxide reductase-Nos) (Zumft, 1997). The last step in the process, N_2_O reduction, is an important focus in denitrification and greenhouse gas research (Jones *et al.*, 2014; Liu *et al.*, 2014; Richardson *et al.*, 2009) as it determines whether the final gaseous product of denitrification is the greenhouse gas N_2_O or the harmless inert gas N_2_. In fact, N_2_O reductase is the only known biological sink of N_2_O (Thomson *et al.*, 2012), therefore, encouraging complete denitrification at the time of N_2_O production represents an important strategy to preventing further rise in atmospheric concentrations (Richardson *et al.*, 2009). In reality, N_2_O vs. N_2_ production is not binary (only N_2_O or N_2_ produced) and N_2_O to N_2_ product ratios depend on a great number of factors including pH (Simek and Cooper, 2002), carbon and nitrate (NO_3_^-^) availability (Senbayram *et al.*, 2012), as well as nitrite (NO_2_^-^) (Firestone *et al.*, 1979; Gaskell *et al.*, 1981).

Conceptually, factors affecting N_2_O emission ratios can be separated into i) microbial community genetic potential for each denitrification step, ii) distal factors, determining that genetic potential in the long term, and iii) proximal factors, acting within genetic potential on short term time scales to impact instantaneous denitrification rates (e.g. carbon and NO_3_^-^ concentrations), (Wallenstein *et al.*, 2006; Groffman *et al.*, 1988). There has been some debate over the relative importance of these factors and disentangling their effects can be difficult when factors such as pH have both immediate effects on enzymatic activity during denitrification and distal effects on denitrification potential (Samad *et al.*, 2016b).

The effect of pH on soil N_2_O emissions is well documented (Simek and Cooper, 2002). Low soil pH results in higher soil N_2_O/(N_2_O+N_2_) ratios, most clearly demonstrated in pH manipulations of soils from the same site (Čuhel *et al.*, 2010; Liu *et al.*, 2010; Simek and Cooper, 2002), but also manifests in differences in N_2_O product ratios between sites (Samad *et al.*, 2016b). Bergaust *et al.* (2010) showed evidence for a post-transcriptional effect on the formation of functional N_2_O reductase at low pH in pure culture experiments, possibly due to impeded assembly of this periplasmic enzyme at low pH. A similar post transcriptional phenomenon was supported using microbial consortia extracted from soils with different native pH (Liu *et al.*, 2014). Contrastingly, some studies have demonstrated that pH effects on N_2_O reduction are dependent on the concentration of NO_3_^-^ or NO_2_^-^ (Blackmer and Bremner, 1978; Firestone *et al.*, 1979; Gaskell *et al.*, 1981). Blackmer and Bremner (1978) showed that pH had negligible effects on N_2_O reduction activity of soils in the absence of supplied NO_3_^-^ while other studies observed and increased inhibitory impact of NO_3_^-^ or NO_2_^-^ under decreasing pH (Firestone *et al.*, 1979; Gaskell *et al.*, 1981). To confuse matters further, pH probably also has long term distal effects on denitrification potential due to its well-known impact on microbial community structuring (Lauber *et al.*, 2008; Kaminsky *et al.*, 2017) and probably more specifically the abundance and ratios of denitrification genes (e.g. Samad *et al.*, 2016b; Domeignoz-Horta *et al.*, 2015; Jones *et al.*, 2014).

The functional and taxonomic composition of denitrifier communities has become an important focus of denitrification research due to advances in molecular tools. Denitrifying microbes may carry all or only some of the full denitrification gene repertoire and therefore changes in the phylogenetic composition of denitrifier communities can affect the ratio of genes coding for N_2_O reductase to those coding for N_2_O producing enzymes, thus determining the genetic potential for N_2_O emission (Graf *et al.*, 2014; Roco *et al.*, 2017). Graf *et al.* (2014) showed that the organisms carrying the *nosZII* gene encoding nitrous oxide reductase clade II commonly had a truncated denitrification pathway without the genes encoding the preceding denitrification steps. Implicit to this finding is the suggestion they may reduce N_2_O/(N_2_O+N_2_) ratios by acting as N_2_O sinks. Jones *et al.* (2014) showed evidence for N_2_O sink capacity related to low *nosZI/nosZII* ratios, though controversial because the results could also be explained as a direct (proximal) effect of soil pH (Bakken *et al.*, 2015). N_2_O/(N_2_O+N_2_) product ratios have been linked to differences in overall microbial community structure as measured by 16S rRNA gene sequencing, suggesting that this could be used as a predictor of N_2_O emission potential (Morales *et al.*, 2014; Samad *et al.*, 2016b). However, it remains unclear whether these correlations indicate a true causal relationship. For example Samad *et al.* (2016b) linked 16S community composition to soil N_2_O/(N_2_O+N_2_) product ratios but both these measures were also correlated to soil pH. Therefore, the results are possibly explained by the well documented but separate effects of pH on N_2_O/(N_2_O+N_2_) (Simek and Cooper, 2002) and on microbial community composition (Lauber *et al.*, 2009; Kaminsky *et al.*, 2017).

N_2_O accumulation during denitrification may also be caused by differential flow of electrons to the separate N-reductases (Pan *et al.*, 2013). In wastewater treatment, low carbon (reductant) availability enhances competition for electrons between N_2_O and upstream N-reductases which can result in transient N_2_O accumulation (Pan *et al.*, 2013; Ribera-Guardia *et al.*, 2014). Indeed carbon and substrate availability can affect N_2_O/(N_2_O+N_2_) product ratios in many contexts including soils, though the direction of the effect is not always consistent (Gillam *et al.*, 2008; Senbayram *et al.*, 2012; Weier *et al.*, 1993).

Here, we aimed to re-assess the consistency of previously outlined (Samad *et al.*, 2016b, 2016a) linkages between N_2_O/(N_2_O+N_2_), microbial community composition (as measured by 16S rRNA gene sequencing and qPCR of *nosZ* genes) and pH in a larger alternative cohort (20 soils) of pasture soils. Previous investigations indicated community composition in the present soil set was correlated to changes in long term rainfall above pH and we hypothesized microbiome to N_2_O emission potential associations would be maintained across this alternate distal driver. Anoxic soil incubations revealed contrasting N_2_O/(N_2_O+N_2_) and denitrification phenotypes based on the timing of N_2_O reduction which determined the propensity for soil N_2_O emission. We further assessed the phenotypes and N_2_O/(N_2_O+N_2_) ratios in relation to potential proximal controls (NO_3_^-^ + NO_2_^-^ concentration, carbon availability) based on 2 alternate hypotheses: 1) N_2_O reduction activity was impaired by higher NO_3_^-^ + NO_2_^-^ concentrations. 2) Impaired N_2_O reduction in high N_2_O/(N_2_O+N_2_) ratio soils is caused by limited carbon availability and thus electron supply to N_2_O reductase.

## Materials and methods

### 2.1 Soil collection

Soils were sampled at 20 sites representing sheep, dairy, beef and goat farms across multiple regions of New Zealand’s South Island. Sampling begun on the 2^nd^ of September 2016 and continued through until the 8^th^ of September. At each site, soil cores (10 cm depth, 2.5 cm diameter) were sampled at 2.5 m intervals across a 7.5 m transect (0 m, 2.5 m, 5 m, 7.5 m) using a stainless steel auger. Triplicate cores were collected for each distance and composited in a bag (12 pooled cores) while a fourth core from each distance was kept separate for molecular analyses (4 cores). If topsoil was less than 10cm deep, additional cores were taken to make up the volume. Soil cores were stored in partially open ziplock bags (to prevent anoxia) on ice until sampling was completed. Pooled cores were homogenized, and worms, insects, grass and large roots removed before field moist storage at 4°C. Core samples for molecular analysis were immediately frozen at −80°C until DNA extraction. See Table S1 for basic soil descriptors based on current analyses and data collected from separate sampling in 2011 (Wakelin *et al.*, 2013).

Pooled site cores were transported to the Norwegian University of Life Sciences (NMBU, Ås, Akershus, Norway), where they were sieved (2mm) and stored at 4°C before initiating kinetic experiments.

### 2.2 Nitrate + nitrite measurements

Endogenous soil nitrate + nitrite (NO_3_^-^ + NO_2_^-^) content was measured in sieved pooled soils using 0.2g of each soil with 1mL of 2M KCl extractant in 1.5 mL microfuge tubes (Lim *et al.*, 2018). Slurries were shaken and spun down at 16000G for 2 minutes before recovering the supernatant into fresh 1.5mL microfuge tubes. NO_3_^-^ + NO_2_^-^ concentration was quantified by chemical reduction to nitric oxide (NO) followed by chemiluminescent detection as detailed by (Braman and Hendrix, 1989;Lim *et al.*, 2018). In brief, 10µL of supernatant was introduced into a sealed glass piping system containing a heated (95°C) acid vanadium chloride solution (50mM VCl_3_ in 1M HCl). VCl_3_ reacts to reduce NO_3_^-^ quantitatively to NO_2_^-^ before converting it to NO gas. The NO gas was captured and carried in an N_2_ stream to a Sievers Nitric Oxide Analyzer 280i system (GE Analytical Instruments, Boulder, CO, USA) for quantification. Standard KNO_3_ solutions (100 to 0.01mM) were used to calibrate the area under detection signal peaks allowing calculation of NO_2_^-^+NO_3_^-^ concentration in soil supernatants. NO_2_^-^+NO_3_^-^ per gram soil or per µL porewater was back calculated based on KCl dilution factor, and soil dry weight/moisture weight (as determined gravimetrically below).

### 2.3 Nitrate adjustment

Soil moisture was determined gravimetrically by drying 5-10g soil at 60°C for minimum 24h (ASTM D2216-10, 2010). Prior to incubation, soils were supplemented with NO_3_^-^ and ammonium by flooding and draining with a 2mM NH_4_NO_3_ solution e.g. (Samad *et al.*, 2016a; Liu *et al.*, 2010; Qu *et al.*, 2014). NO_3_^-^ supplied N for denitrification while ammonium acted as a preferential assimilatory N source. For this, an 80g dry weight equivalent of soil was placed in a 500mL Sterafil Filter Holder (Merck, Burlington, MA, USA) and flooded with 300mL of 2mM NH_4_NO_3_ (sufficient volume to dilute endogenous NO_3_^-^). After 15min the solution was drained through a 0.2µM cellulose filter (Merck) with 1.2µM glass-fibre pre filter (Merck) using a vacuum manifold. Soils were mixed and a subsample was taken for overnight moisture content analysis (5g, as above). The remainder of the soils was stored overnight in funnels covered with aluminum foil before use in incubation experiments the next day.

### 2.4 Gas kinetics measurements

Respiration and denitrification activity of soils was measured by gas chromatography of headspace gases (O_2_, CO_2_, NO, N_2_O, N_2_) in oxic and anoxic batch incubations commencing between 39 and 56 days post sampling. The temperature controlled robotic autosampler, gas chromatograph (Agilent GC −7890A equipped with ECD, TCD, FID) and chemiluminescence NOx analyzer (Model 200A, Advanced Pollution Instrumentation, San Diego, USA) used here are described in detail by (Molstad *et al.*, 2007; Qu *et al.*, 2014). The device holds up to 44 sealed 120 mL serum vials in a temperature controlled water bath. A robotic arm equipped with a hypodermic needle and a peristaltic pump takes headspace gas samples periodically and pumps them through dedicated sample loops in the GC. The GC uses helium as carrier gas while subsequent back pumping replaces sampled gas with helium thus maintaining the pressure in the serum vials at ∼1 atm. Dilution and leakage are back calculated following experiment completion to allow estimation of true gas production. Here, 20g dry weight equivalent of NH_4_NO_3_ adjusted soil was placed in triplicate 120mL serum vials and crimp sealed with butyl rubber septa. A 15g subsample of each soil was frozen at the start of each incubation for subsequent measurement of NO_2_^-^ +NO_3_^-^ concentrations as described above. The serum vials were placed in the water bath (20°C) under the autosampler and allowed to equilibrate before releasing overpressure through a water filled syringe without piston. Additional vials (duplicates) were filled with premixed standard gases (including 400ppm CO_2_, 10,000ppm CO_2_, 500ppb N_2_O, 151ppm N_2_O, 25ppm NO) supplied by AGA industrial gases (Oslo, Akershus, Norway), compressed air (781,000ppm N_2_, 200900ppm O_2_) or helium. The autosampler was programmed to take headspace samples every ∼5hrs. After 7 rounds of sampling, at ∼40hrs, anoxia was induced by helium flushing the vials (3 cycles of evacuation for 180sec and He-filling for 20sec) and incubation was continued until most soils had converted all denitrification products to N_2_. Two separate experiments were required 10 days apart to process all 20 soils.

### 2.5 Measures of N_2_O emission potential/kinetics

The N_2_O hypothetically emitted (%) metric was calculated as max µmol N_2_O-N accumulated in vial expressed as a percentage of final cumulative N (N_2_-N). N_2_O hypothetically emitted (%) was used to evaluate the sequentiality of N_2_O production and reduction steps over the course of an incubation and the relative N_2_O emission potential of each soil. It is equivalent to typical N_2_O product ratios (N_2_O/N_2_O +N_2_) but expressed relative to total final accumulated N rather than just gaseous N_2_O+N_2_ at the timepoint of calculation. It is our preferred metric because:

1. It is event based (i.e. calculated at peak in vial N_2_O). This allows comparison of soils with divergent denitrification timescales/rates.
2. It is a relative measure (i.e. expressed as a percentage of total N finally accumulated). This allows comparison of soils with contrasting initial NO_3_^-^ supply and net denitrification rates.
3. It describes N_2_O emission potential (i.e. soils accumulating greater peak N_2_O in headspace are likely to be higher emitters in situ). This N_2_O is likely to be emitted in an unsealed environment therefore this metric ignores net N_2_O reconsumption from the headspace.
4. It directly describes the sequentiality of N_2_O production/reduction (i.e. to what degree N_2_O production and reduction to N_2_ were carried out at the same time and to the same magnitude, thus mitigating N_2_O accumulation). This was highly relevant to the soil kinetics observed here (see results).

Additional emission potential metrics were evaluated to allow comparison with previous studies. N_2_O/(N_2_O+N_2_) (N_2_O ratio) calculated as µmol cumulative N_2_O-N (per vial) over max µmol cumulative N_2_O-N plus µmol cumulative N_2_-N at various timepoints (N_2_O/(N_2_O+N_2_) (max N_2_O), N_2_O/(N_2_O+N_2_) (50hrs)) (Samad *et al.*, 2016a). N_2_O index (I_N2O_) as used previously (Liu *et al.*, 2010; Qu *et al.*, 2014; Samad *et al.*, 2016a), calculated as the area under a N_2_O curve over the area under an N_2_O curve + N_2_ curve using the following formula: 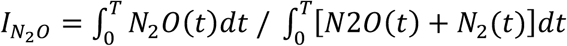

It is useful because it allows a time-integrated view of N_2_O vs. N_2_ stoichiometry. Areas are calculated for each time period between two sampling points (∼5hrs) and summed up to an arbitrary time point (T). Here, we used 50hrs (I_N2O_ (50hrs)) and once all denitrification gas was accumulated as N_2_ (I_N2O_ (N_2_ plateau)).

N_2_O hypothetically emitted (%) value cut-offs were assigned to separate N-gas accumulation patterns into discrete phenotypes based on N_2_O, NO, and N_2_ accumulation patterns. Sequential soils (88% to 100%) had close to 0 accumulation of N_2_ before peak N_2_O, a sudden increase in N_2_ rate was observed after peak N_2_O, and a single often high NO peak was usually observed. Concurrent soils (0 to 80%) developed N_2_ production early which was either sustained or gradually increased. NO levels were typically controlled after a brief peak. Intermediate soils (81% to 87%) had features of both phenotypes. NO was typically poorly controlled, some N_2_ production was observed before peak N2O but sharp increases in N_2_ rate were observed after peak N_2_O.

### 2.6 pH measurement

For each soil, a 10mL subsample was placed in a plastic container using a volumetric spoon and 20mL of 10mM CaCl_2_ was added. Containers were capped and shaken until the soil was dispersed in the solution. Soils were left overnight at room temperature. Soils were re-dispersed by shaking and left to settle for 10min before the pH was measured in the supernatant using a H170 pH meter (Hach, Loveland, CO, USA).

### 2.7 Rainfall

Average daily rainfall at the sample sites (mm day^-1^) was estimated at various timescales (month, year, 10 years) using rainfall data from New Zealand’s national climate database. Data were assessed through the CliFlo web system (NIWA, 2017). Collected data spanned from 08/09/96 to 07/09/2016. We also included rainfall and potential evapotranspiration estimates from a previous study (Wakelin *et al.*, 2013) on the same soils (rainfall historical) which used average daily measurements for 5 years prior to sampling. Values were calculated through interpolations (Cichota *et al.*, 2008; Tait *et al.*, 2005) using the Virtual Climate Station from NIWA (Wellington, Wellington, New Zealand). Drainage class (1: very poor to 5: well) was collected from the New Zealand Fundamental soil Layer (LRIS, 2020).

### 2.8 DNA extraction

Distance-specific site cores (0m, 2.5m, 5m, 7.5m) were defrosted for DNA extraction. To test if pooling and sieving was necessary for future sampling, a subsample of 15g fresh weight from each core was pooled into a mixed sample (mixed-m) and a subsample of this sample was sieved through a 2mm sieve (mixed and sieved-ms). ∼0.25g of each distance specific sample per site and the two additional m and ms samples per site (6 samples per site times 20 sites = 120 samples) were extracted using a Powersoil DNA Isolation kit (Mobio, Carlsbad, CA, USA) according to the standard instructions. Bead beating was carried out at 1500rpm in two 15sec steps with intermittent cooling using a 1600 MiniG cell-lyser (SPEXSamplePrep, Metuchen, NJ, USA). DNA extracts were quantified and quality checked using a Qubit fluorometer (Invitrogen, Carlsbad, CA, USA) with Qubit dsDNA HS assay (Invitrogen) and NanoDrop One (Thermo Scientific, Waltham, Massachusetts, USA). pH-CaCl_2_ and pH-H_2_O of ms soils was measured as described earlier (section 2.2.2, pH measurement) but using a MP22O pH meter with Inlab 413 electrode (Mettler Toledo, Columbus, Ohio, USA).

### 2.9 16S amplicon sequencing

16S amplicon sequencing was carried out on a single lane of an Illumina HiSeq using Version 4_13 of the Earth Microbiome Project standard protocol (Caporaso *et al.*, 2012). Open reference OTU picking (97% similarity, UCLUST (Edgar, 2010) and taxonomy assignment (BLAST (Altschul *et al.*, 1990) was carried out in QIIME 1.9.1 (Caporaso *et al.*, 2010) using version 128 of the SILVA database (Quast *et al.*, 2013). Site-specific sequence pools were then subsampled 10 times to a depth of 37120 sequences. Subsampled pools were averaged using basic R functions (R Core Team, 2016). NMDS ordinations (Bray Curtis dissimilarity) were carried out using Phyloseq (McMurdie and Holmes, 2013). Mantel tests were carried out in Vegan (Dixon, 2003) using a Pearson correlation method. Sequences have been submitted to the NCBI Sequence Read Archive under accession numbers SRR11650167 to SRR11650286 and BioProject ID PRJNA629050.

### 2.10 qPCR

Total prokaryotic abundance, and nitrous oxide reductase gene abundance for Clade I and II were measured by targeting the 16S rRNA gene and *nosZ* gene respectively, using the following primer pairs: 16S UNIV F&R (Hartman *et al.*, 2009), nosZ2F & nosZ2R (Henry *et al.*, 2006), 1153_nosZ8F & 1888_nosZ29R (Jones *et al.*, 2013). Reactions (10µL total volume) consisted of 10ng soil DNA, forward and reverse primers at a final concentration of 0.5µM (except for *nosZ II* reactions which included 1µM), 5µL of Luminaris HiGreen low Rox qPCR Master Mix (Thermo Scientific) and nuclease free water (Thermo Scientific) to make up the 10µL volume. Minimum triplicate reactions per sample were performed using a QuantStudio 6 flex qPCR machine (Applied Biosystems, Foster City, CA, USA) according to the following thermal cycling conditions. 16S: 2 min UDG pre-treatment at 50°C, 10min initial denaturation at 95°C, 40 cycles of 15sec denaturation at 95°C, 30sec annealing at 65°C and 30sec extension at 72°C. *nosZI*: 2 min UDG pre-treatment at 50°C, 10min initial denaturation at 95°C, 40 cycles of 15sec denaturation at 95°C, 30sec annealing at 58.5°C and 30sec extension at 72°C. *nosZII* touchdown: 2 min UDG pre-treatment at 50°C, 10min initial denaturation at 95°C, 6 cycles of amplification decreasing annealing temperature by 1°C per cycle consisting of 15sec denaturation at 95°C, 30sec annealing at 60-55°C and 30sec extension at 72°C, 44 cycles of amplification consisting of 15sec denaturation at 95°C, 30sec annealing at 54°C, 30sec extension at 72°C and 30sec at 80°C for signal detection. All reaction plates included minimum triplicate no-template controls and a 10-fold dilution series of pGEM-t-easy (Promega, Madison, WI, USA) cloned standards for the relevant amplicon, encompassing the sample quantification range. Measurement of the desired amplicon was confirmed by a melt curve analyses (15sec denaturation at 95°C, 1min 60°C, 30 sec 95°C) following target amplification.

### 2.11 Predicted NO_3_^-^ + NO_2_^-^ accumulation and predicted nitrification

Predicted NO_3_^-^ + NO_2_^-^ at the start of anoxia was calculated as the final accumulated µmol-N accumulated in each vial at the end of the incubation. Predicted nitrification during the oxic period was calculated as the difference between measured NO_3_^-^ + NO_2_^-^ at the beginning of the experiment and predicted NO_3_^-^ + NO_2_^-^ at the start of anoxia.

### 2.12 Carbon amendment experiment

An independent experiment was set up as described above with some modifications to test the influence of carbon availability on gas kinetics. Incubations commenced 3 months after initial sampling. Five soils were selected based on covering a range of N_2_O ratios (40-Fairlie Geraldine, 20-Waitaha, 1-Woodend, 33-Rae’s Junction, 5-Waipapa). In these incubations, an initial period of oxic storage and incubation was not carried out and the concentration of NH_4_NO_3_ in flooding solutions was increased (4mM) to account for extra NO_3_^-^ accumulated during the oxic period in the original incubations. Incubations were monitored under two treatment conditions: 4mM NH_4_NO_3,_ ± 10mM sodium glutamate as a carbon source. Glutamate can be utilised by most bacteria and in addition could provide a preferential organic N source preventing NO_3_^-^ assimilation (Halvorson, 1972). Sodium glutamate solutions were pH adjusted to the soils’ native pH using HCl.

### 2.13 Statistical analyses

Incubation kinetics variables are presented and used in correlations as the mean of triplicate incubations or duplicate incubations in cases where a replicate had to be dropped due to gas leakage. Spearman’s ranked correlations between incubation kinetic measures and soil variables were used based on non-normaly distributed data. Discrete phenotypic groups were compared to incubation variables using a Wilcoxon rank sum test (differences of medians) with chi-squared approximation (but are also supported with continuous/ranked analyses). NMDS plots were evaluated based on stress <0.2. NMDS plots are presented for a single pooled sample per soil for appropriate statistical comparison to gas kinetic and environmental variables but ordinations with full distance specific replicates are available (Figure S3). Bray Curtis dissimilarity matrixes were compared against incubation and environmental variables by Mantel test using a pearson correlation co-efficient.. NMDS axis 1 and 2 co-ordinates were also extracted and tested against incubation and environmental variables using a spearman’s ranked correlation to identify variables associated with a particular axis. Multiple linear regression for prediction of N_2_O hypothetically emitted (%) using rainfall, soil drainage class and potential evapotranspiration was performed using standard least squares.

## Results

### 3.1 Denitrification gas kinetics

We monitored gas production (CO_2,_ NO, N_2_O, N_2_) from NH_4_NO_3_ amended soil incubations to identify soils with contrasting denitrification gas production kinetics and potential for N_2_O emission (key incubation variables available in Table S2). Soils were initially incubated under oxic conditions (40hrs) to identify their aerobic respiratory potential. Soil CO_2_ production was in the range of 1 to 5µmol hr^-1^ vial^-1^ (mean ± SD = 3.43 ± 2.89) with the exception of one soil (18-Kumara, a flipped pasture) that had a production rate of 15.21µmol hr^-1^ vial^-1^.

During the subsequent anoxic incubation period, soils varied greatly in the timing of N_2_O production and further reduction to N_2_: while some soils carried out concurrent N_2_O production/reduction from the onset of anoxia, others carried out each step sequentially, accumulating most N as N_2_O before converting it stoichiometrically to N_2_ (Figure 1). We evaluated the sequentiality of N_2_O production/reduction on a continuous scale using N_2_O hypothetically emitted (%) and applied somewhat arbitrary cutoffs (see section 2.5.) to place each soil in discrete phenotypic groups: concurrent (0 to 80%), intermediate (81% to 87%) and sequential (88% to 100%). In addition to timing of N_2_O production/reduction, alternative phenotypic groups had contrasting NO accumulation patterns: more concurrent N_2_O producing/reducing soils accumulated far less NO (Spearman’s correlation, average µmol NO vial^-1^ vs. N_2_O hypothetically emitted %, ρ=0.80, p<0.0001), and most displayed a very low pseudo steady state NO level after a brief peak in accumulation (Figure 1A).

**Figure 1.**
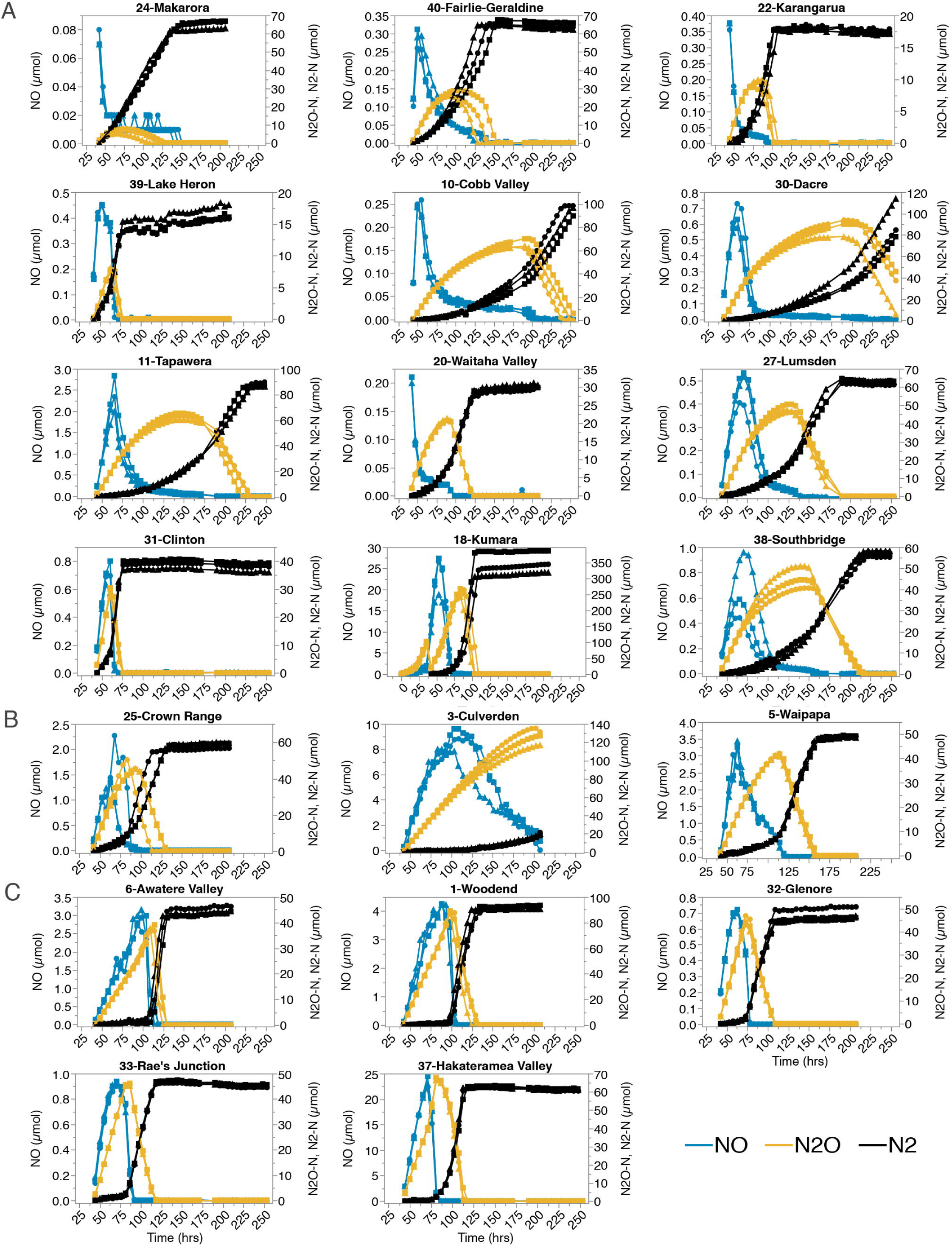
N gas kinetics of 20 NZ pasture soils amended with 2mM NH_4_NO_3_ and incubated in a helium atmosphere. The figure shows the wide variation in timing of N_2_O reduction (N_2_ production) leading to variable N_2_O accumulation. N_2_O hypothetically emitted (%) (Table S2) was used to evaluate sequentiality of N_2_O production/reduction on a continuous scale (soils ordered top left to bottom right) and define soil phenotypes (concurrent (A), intermediate (B) and sequential (C)) based on discrete arbitrary cutoffs. Circles, squares, triangles represent three replicate vials. Concurrent N_2_O production/reduction, is associated with specific NO emission pattern: Lower NO accumulation eventually stabilising at a pseudo steady state. Sequential soils usually accumulate higher max NO. Note that the panels have different scaling of N_2_O (Orange), N_2_ (Black) and NO (Blue) and values are reported as µmol-N per vial. Scaled version available (Figure S1). Version with CO_2_ available (Figure S2).

N_2_O hypothetically emitted (%) was also used to evaluate soil N_2_O emission potential as sequentiality of N_2_O production/reduction determines N_2_O accumulation, while omitting the reconsumption of headspace N_2_O predominant late in sequential soils which is more likely to be emitted in an unsealed environment. Alternative measures of N_2_O emission potential were also evaluated to maintain comparability with previous studies and gave similar soil rankings (Spearmans correlation vs. N_2_O hypothetically emitted %, N_2_O/(N_2_O+N_2_) (max N_2_O): ρ=0.94, p<0.0001, N_2_O/(N_2_O+N_2_) (50hrs): ρ=0.71, p<0.001, I_N2O_ (N_2_ plateau): ρ=0.79, p<0.0001, I_N2O_ (50hrs): ρ=0.53, p<0.05).

### 3.2 Interaction between N_2_O emission potential, pH and community composition

We hypothesized that the observed kinetic patterns/N_2_O emission potentials were linked to differences in pH and community differences based on previously observed linkages between N_2_O product ratios, pH and 16S microbial community composition using the same incubation methodology (Samad *et al.*, 2016b). N_2_O hypothetically emitted (%) was not correlated with soil pH (Spearman’s correlation, measures of N_2_O emission vs. pH (CaCl_2_ or H_2_O), p > 0.05), but did map to differences in 16S community composition across axis 1 in NMDS plots (Table 1, Spearman’s correlation NMDS axis 1, Figure 2A). However, significant correlation to the full dissimilarity matrix based on Mantel tests (Table 1) was not achieved unless all distance specific replications were included in analyses (Figure S3). Similar trends were observed for alternative emission potential metrics (Table 1). pH and long-term average daily rainfall were identified as potential drivers of differences in microbial community composition. Both were significantly correlated to overall changes in the dissimilarity matrix, and mapped primarily to NMDS axis 1 and 2 respectively (Table 1, Figure 2B)

**Table 1.**
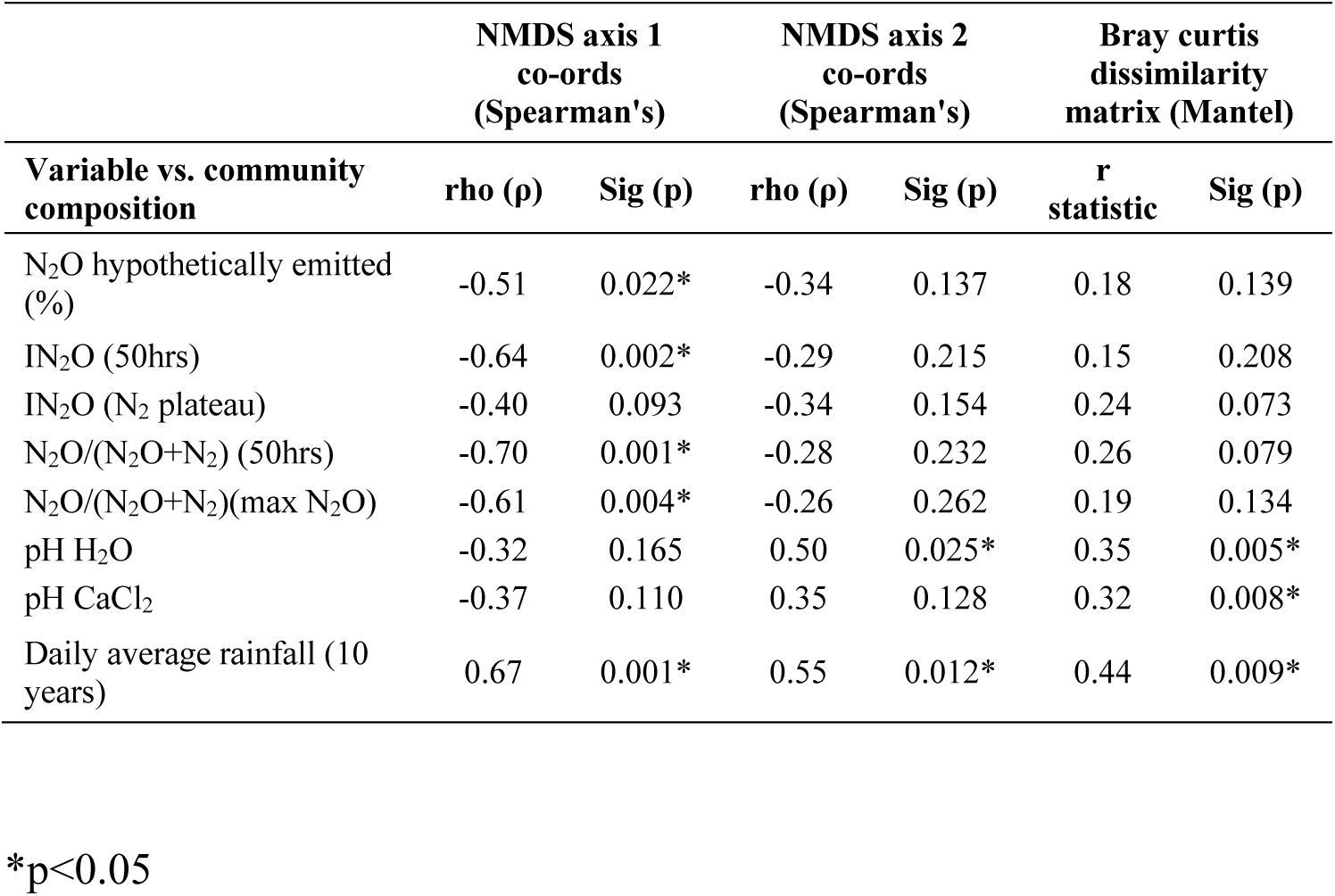
N_2_O emission potential and other variable correlations to community dissimilarity.

| Variable vs. community composition | NMDS axis 1<br>co-ords<br>(Spearman's) |  | NMDS axis 2<br>co-ords<br>(Spearman's) |  | Bray curtis<br>dissimilarity<br>matrix (Mantel) |  |
| --- | --- | --- | --- | --- | --- | --- |
|  | rho (ρ) | Sig (p) | rho (ρ) | Sig (p) | r<br>statistic | Sig (p) |
| N<sub>2</sub>O hypothetically emitted (%) | -0.51 | 0.022* | -0.34 | 0.137 | 0.18 | 0.139 |
| IN<sub>2</sub>O (50hrs) | -0.64 | 0.002* | -0.29 | 0.215 | 0.15 | 0.208 |
| IN<sub>2</sub>O (N<sub>2</sub> plateau) | -0.40 | 0.093 | -0.34 | 0.154 | 0.24 | 0.073 |
| N<sub>2</sub>O/(N<sub>2</sub>O+N<sub>2</sub>) (50hrs) | -0.70 | 0.001* | -0.28 | 0.232 | 0.26 | 0.079 |
| N<sub>2</sub>O/(N<sub>2</sub>O+N<sub>2</sub>)(max N<sub>2</sub>O) | -0.61 | 0.004* | -0.26 | 0.262 | 0.19 | 0.134 |
| pH H<sub>2</sub>O | -0.32 | 0.165 | 0.50 | 0.025* | 0.35 | 0.005* |
| pH CaCl<sub>2</sub> | -0.37 | 0.110 | 0.35 | 0.128 | 0.32 | 0.008* |
| Daily average rainfall (10 years) | 0.67 | 0.001* | 0.55 | 0.012* | 0.44 | 0.009* |

\*p<0.05

**Figure 2.**
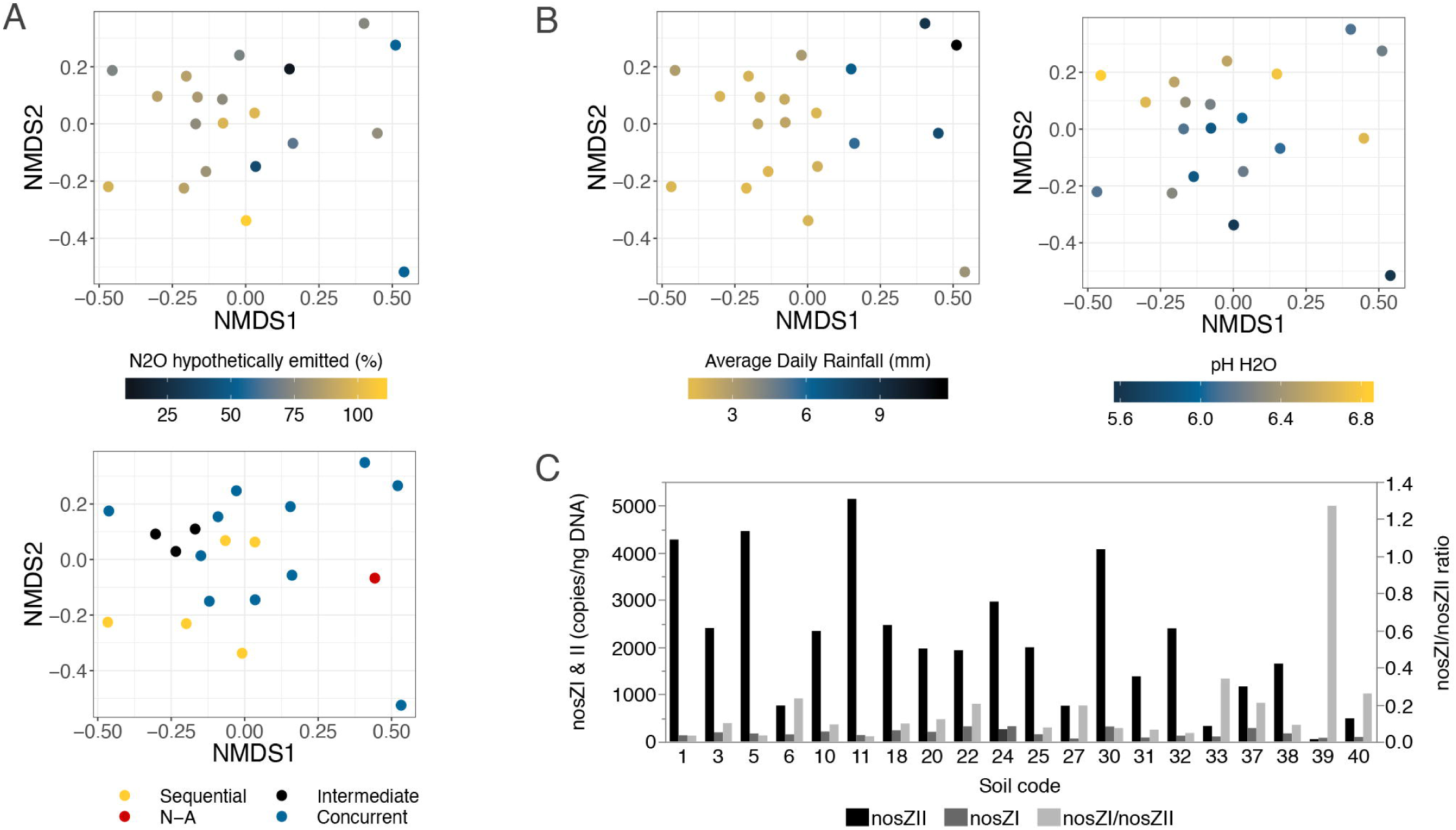
Microbial community analyses reveal links between 16S community composition and N_2_O emission potential/phenotypes (A), average daily rainfall over 10 years and pH (B). qPCR reveals greater abundance of nosZII relative to nosZI (C). NMDS ordination plots (A, B) compare prokaryotic dissimilarities (Bray Curtis) of a single pooled mixed sieved soil sample per site. Full distance specific replications are presented in Figure S3. Stress values for ordinations were 0.14. Correlations between variables and NMDS axes or the Bray Curtis dissimilarity matrix are presented in Table 1.

We also measured *nosZII* gene copy numbers as a community related functional metric as it has been suggested *nosZII* carrying organisms are important for soil N_2_O reduction activity. *nosZII* copy number expressed in various forms (numbers per ng soil DNA, per gram of soil, normalized to 16S copy numbers and relative to *nosZI* copy numbers) were not significantly correlated to N_2_O hypothetically emitted % or other measures of emission potential (p>0.05, Spearman’s correlation). *nosZII* copy numbers were most strongly correlated with soil pH CaCl_2_ (Spearman’s correlation, ρ=0.60, p<0.01) and interestingly, were ∼10 fold higher in abundance than *nosZI* (Figure 2C).

### 3.3 Rainfall

In addition to community associations, we assessed the direct relationship between rainfall and soil phenotypes/N_2_O emission potential. More concurrent N_2_O production/reduction was associated with higher long-term average daily rainfall (Figure 3A). The strength of the correlation was best (and highly significant; p<0.01) when rainfall regime was averaged over a prior year or decade, and was not significant when averaged over shorter time span (Table S3). Linear regression of rainfall (10 years) and hypothetically emitted N_2_O (%) poorly recapitulated the trends observed in non-parametric and non-continuous analyses (Figure 3B), probably due to the high variability in average daily rainfall among low N_2_O emitting soils. Correlations between all rainfall and N_2_O emission potential metrics are found in Table S3. Multiple linear regression indicated drainage class and potential evapotranspiration did not aid prediction of N_2_O hypothetically emitted (%) with only rainfall producing a significant parameter effect (p = 0.03, Table S4).

**Figure 3.**
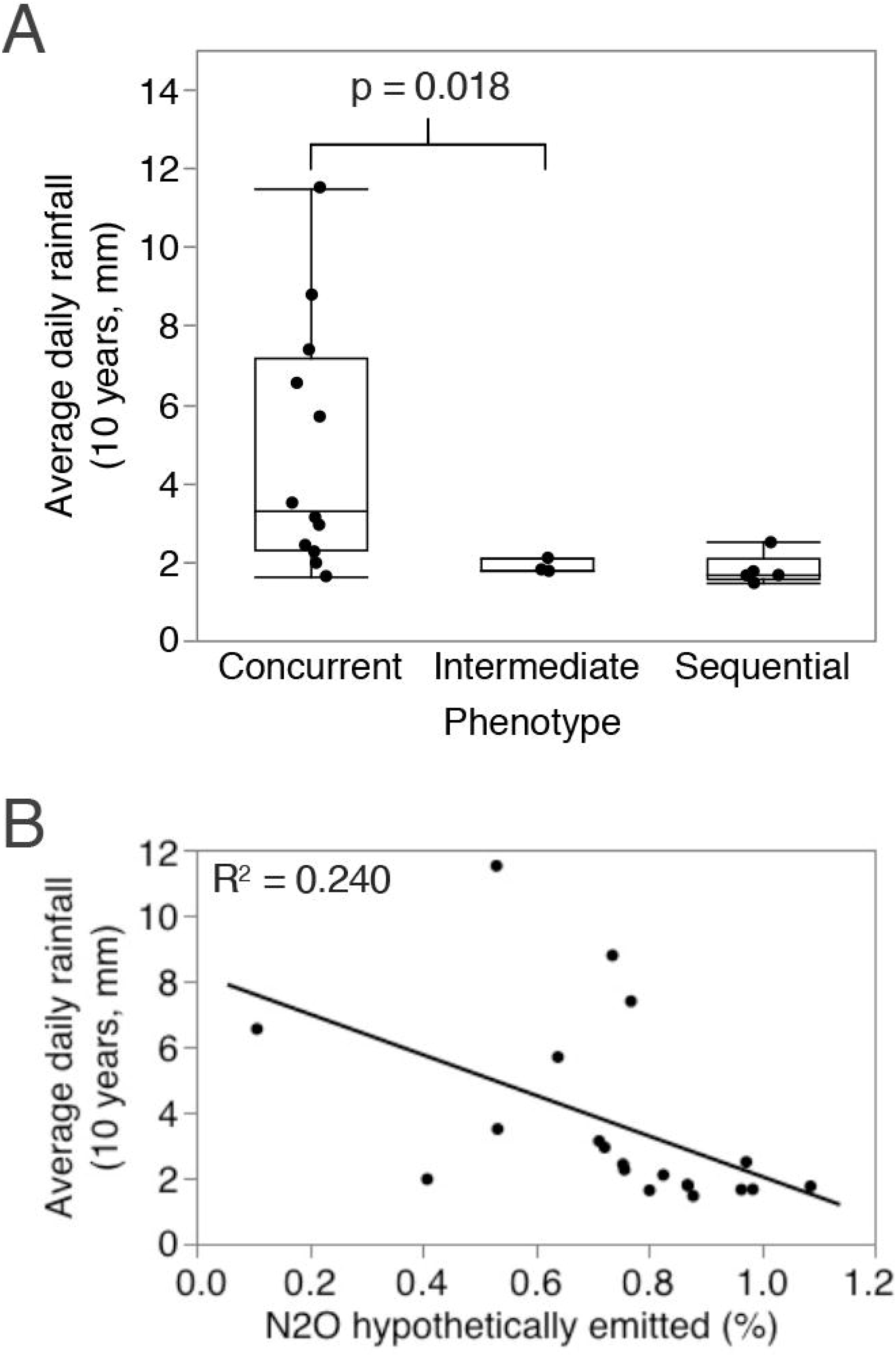
Relationship between average daily rainfall (average daily mm rainfall over 10 years prior to sampling) and N_2_O production/reduction phenotypes (A) or N_2_O hypothetically emitted % (B). p-value presented is for difference of medians using Wilcoxon rank sum test with chi-squared approximation. Correlations between all rainfall and N_2_O emission potential metrics are presented in **Table S3**.

### 3.4 Nitrate + nitrite

Final cumulated N_2_ levels per vial were inconsistent between soils suggesting significant variation in NO_3_^-^ + NO_2_^-^ concentrations upon initiation of the anoxic incubation period (Figure 4A). Further, comparison of measured soil NO_3_^-^ + NO_2_^-^ before incubation and at the start of the anoxic incubation estimated from cumulative denitrified N (Figure 4A) suggested NO_3_^-^ or NO_2_^-^ was accumulated during the oxic incubation period, presumably due to nitrification of added ammonium. We investigated the potential impact of these variable initial NO_3_^-^ + NO_2_^-^ concentrations as it has previously been demonstrated that N_2_O reduction activity is sensitive to NO_3_^-^ and NO_2_^-^ concentration. Predicted NO_3_^-^ + NO_2_^-^ porewater concentrations at the start of the anoxic period were not significantly correlated to N_2_O hypothetically emitted %, however, significant positive correlations were observed for some alternative measures of N_2_O emission potential (Table 2).

**Figure 4.**
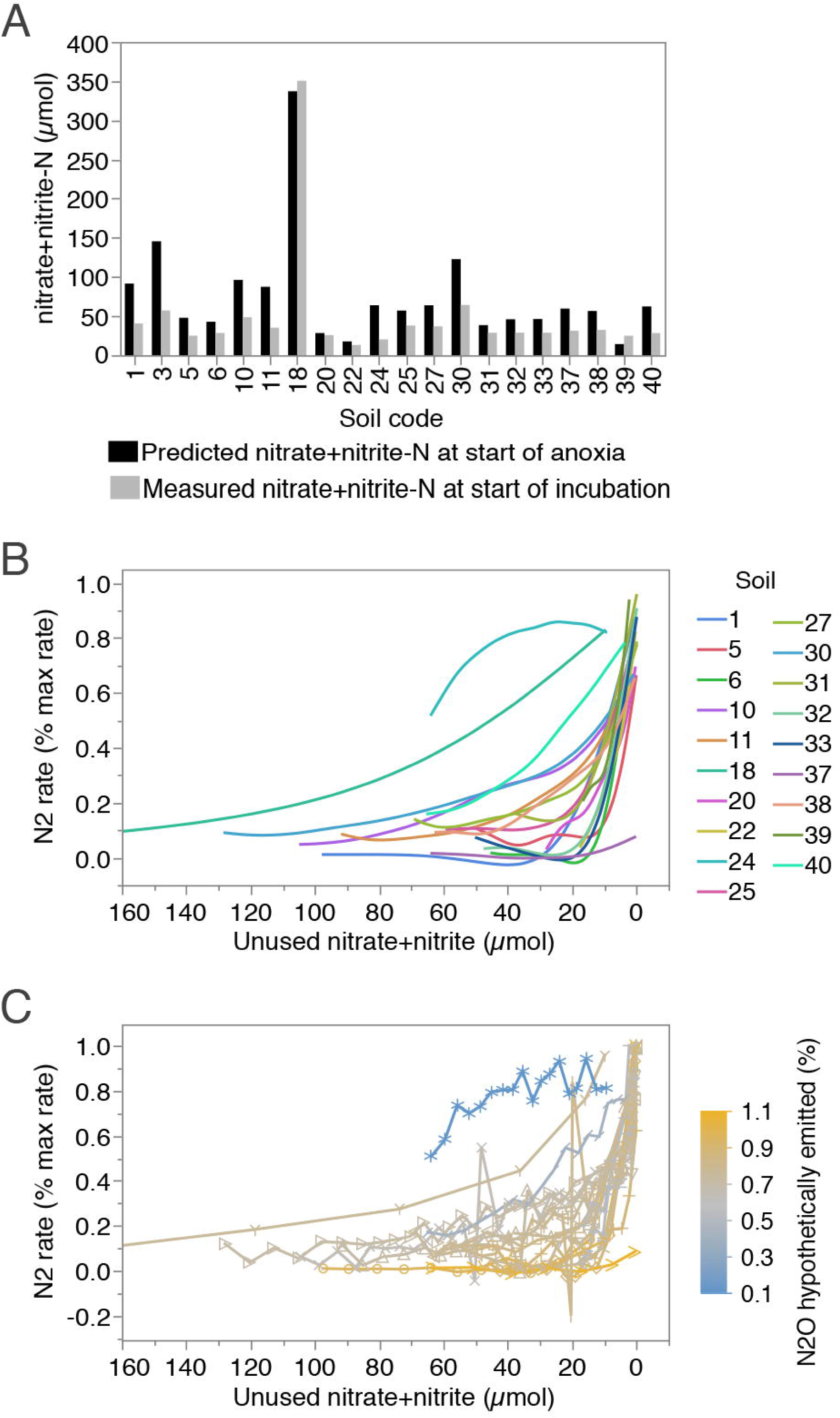
Normalized soil N_2_ production rates (N_2_ rate over max N_2_ rate in same soil) increase as available NO_3_^-^ + NO_2_^-^ is depleted in anoxic soil incubations amended with 2mM NH_4_NO_3_. In most soils, predicted NO_3_^-^ + NO_2_^-^ at beginning of anoxic period is greater than measured NO_3_^-^ + NO_2_^-^ immediately following NO_3_^-^ amendment (A) indicating NO_3_^-^ + NO_2_^-^ accumulation, most likely due to nitrification during oxic pre-incubations. Plots (C, D) allow comparison of N_2_O reduction activity at similar NO_3_^-^ + NO_2_^-^ concentration for each soil. Ranking of soils across Y-axis could indicate potential variation in soil N_2_O reduction sensitivity to NO_3_^-^ + NO_2_^-^ concentration, however this ranking simply describes the aforementioned N_2_O production/reduction phenotypes (C) and a potential effect cannot be separated from the natural progression of denitrification or other biases.

**Table 2.**
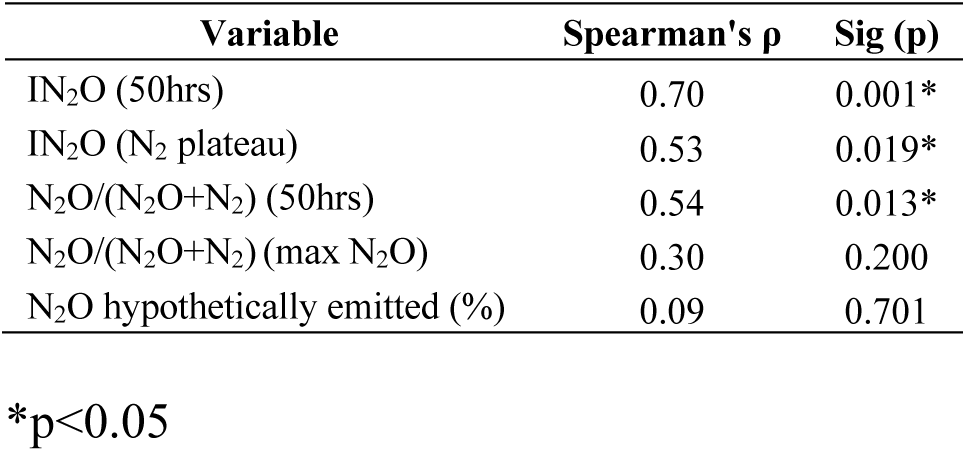
Correlations between predicted NO_3_^-^ + NO_2_^-^ at the start of anoxia and measures of N_2_O emission potential.

| Variable | Spearman's ρ | Sig (p) |
| --- | --- | --- |
| IN<sub>2</sub>O (50hrs) | 0.70 | 0.001* |
| IN<sub>2</sub>O (N<sub>2</sub> plateau) | 0.53 | 0.019* |
| N<sub>2</sub>O/(N<sub>2</sub>O+N<sub>2</sub>) (50hrs) | 0.54 | 0.013* |
| N<sub>2</sub>O/(N<sub>2</sub>O+N<sub>2</sub>) (max N<sub>2</sub>O) | 0.30 | 0.200 |
| N<sub>2</sub>O hypothetically emitted (%) | 0.09 | 0.701 |

\*p<0.05

Normalized N_2_ production rates (% of maximum) were plotted against residual NO_3_^-^ + NO_2_^-^ concentrations estimated from denitrification progress at different stages during the anoxic incubation to allow comparison of soil N_2_O reduction rates at similar NO_3_^-^ + NO_2_^-^ concentrations (Figure 4B, C). Concurrent soils showed greater % N_2_ production rate at greater NO_3_^-^ + NO_2_^-^ levels while most sequential phenotype soils maintained near zero N_2_ production rates until NO_3_^-^ + NO_2_^-^ fell below 20-10µmol (Figure 4C). However, it should not necessarily be concluded that different sensitivities of N_2_O reduction to NO_3_^-^ + NO_2_^-^ explain the variation in soil N_2_O reduction/production phenotype, due to confounding by time, N denitrified, the natural progression of denitrification and other unknown factors. Sensitivity of N_2_O reduction to NO_3_^-^ + NO_2_^-^ has previously been explained by pH, therefore we overlayed pH onto plots but did not find an explanatory pattern (Figure S4).

### 3.5 Carbon supplementation

We hypothesized that differences in apparent denitrification phenotypes resulted from electron competition under carbon-limited conditions between earlier steps of denitrification and N_2_O reduction. In further incubations, soils representing a range of phenotypes/N_2_O emission potentials were amended with both glutamate (to relieve potential carbon limitation) and NH_4_NO_3_ (to provide NO_3_^-^ for denitrification) or control vials with NH_4_NO_3_ alone. In most cases, carbon negative controls (Figure 5, left panel) recapitulated the general phenotypic trends observed in initial incubations (Figure 1) but there were some large observable differences, possibly caused by changes to the incubation preparation methodology (omitted oxic preincubation, increased added NH_4_NO_3_ to 4mM). In particular, soil 40-Fairlie-Geraldine (Figure 5A, top-left) showed a “weakened” concurrent phenotype, compared with original incubations (N_2_O hypothetically emitted %, 0.41 in original vs. 0.80 in second incubation). Differences for other soils were much less dramatic (Table 3). All +N treatments also had higher CO_2_ production rates and denitrification process rates, on average 1.44 (average CO_2_ production rate) and 1.38 (max N_2_O production rate) times higher respectively, compared with original incubations. This may suggest changes made to incubation methodology resulted in higher respiration and probably more available soil carbon.

**Table 3.** Comparison of gas kinetics features across all incubation treatments for five soils.

| Soil | Concurrent |  |  |  |  |  | Sequential |  |  |  |  |  | Intermediate |  |  |
| --- | --- | --- | --- | --- | --- | --- | --- | --- | --- | --- | --- | --- | --- | --- | --- |
|  | Fairlie-Ger (40) |  |  | Waitaha (20) |  |  | Rae's Junct (33) |  |  | Woodend (1) |  |  | Waipapa (5) |  |  |
|  | 1st | +N | +C+N | 1st | +N | +C+N | 1st | +N | +C+N | 1st | +N | +C+N | 1st | +N | +C+N |
| Av N<sub>2</sub> rate before peak N<sub>2</sub>O (μmol/hr) | 0.38 | 0.36 | 0.10 | 0.17 | 0.14 | 0.06 | 0.16 | 0.08 | 0.10 | 0.08 | 0.18 | 0.22 | 0.11 | 0.26 | 0.54 |
| Av N<sub>2</sub> rate after peak N<sub>2</sub>O (μmol/hr) | 0.98 | 2.20 | 3.08 | 0.58 | 0.52 | 1.87 | 1.26 | 1.57 | 4.65 | 2.62 | 2.77 | 4.22 | 0.84 | 1.30 | 2.40 |
| Max N<sub>2</sub> rate (μmol/hr) | 1.39 | 2.54 | 4.26 | 0.76 | 0.64 | 3.84 | 1.51 | 2.10 | 6.89 | 3.65 | 3.53 | 5.01 | 1.04 | 1.50 | 3.14 |
| Max N<sub>2</sub>O rate (μmol/hr) | 0.91 | 1.60 | 2.46 | 0.71 | 0.84 | 1.15 | 1.25 | 1.59 | 2.77 | 2.24 | 2.81 | 2.83 | 0.79 | 1.11 | 1.87 |
| Estimated Av total N turnover rate (μmol NO<sub>2</sub><sup>-</sup>, NO, N<sub>2</sub>O, N<sub>2</sub> N/hr) | 2.78 | 4.40 | 5.46 | 1.42 | 1.50 | 3.29 | 2.73 | 3.27 | 6.09 | 4.44 | 5.59 | 6.90 | 1.77 | 3.10 | 4.99 |
| Av CO<sub>2</sub> rate (μmol/hr) | 1.15 | 1.92 | 2.68 | 0.90 | 0.92 | 1.64 | 1.34 | 1.51 | 2.55 | 1.99 | 2.91 | 3.61 | 0.74 | 1.27 | 2.28 |
| Av NO (μmol) | 0.10 | 0.67 | 1.15 | 0.04 | 0.04 | 0.56 | 0.61 | 0.63 | 0.62 | 1.54 | 0.27 | 0.23 | 1.34 | 1.50 | 0.51 |
| Max NO (μmol) | 0.29 | 2.21 | 4.64 | 0.21 | 0.27 | 1.27 | 0.92 | 2.31 | 2.51 | 4.25 | 1.53 | 1.23 | 3.25 | 6.06 | 2.79 |
| N<sub>2</sub>OI (N<sub>2</sub> plateau) | 0.44 | 0.61 | 0.69 | 0.57 | 0.58 | 0.80 | 0.65 | 0.65 | 0.69 | 0.66 | 0.63 | 0.70 | 0.64 | 0.53 | 0.55 |
| N<sub>2</sub>O/(N<sub>2</sub>O+N<sub>2</sub>)(max N<sub>2</sub>O) | 0.59 | 0.79 | 0.95 | 0.71 | 0.80 | 0.97 | 0.87 | 0.94 | 0.95 | 0.96 | 0.93 | 0.92 | 0.84 | 0.79 | 0.71 |
| Hypothetically emitted N<sub>2</sub>O (%) | 0.41 | 0.80 | 0.92 | 0.74 | 0.83 | 1.00 | 0.98 | 0.99 | 1.00 | 0.96 | 0.90 | 0.96 | 0.87 | 0.70 | 0.74 |

**Figure 5.**
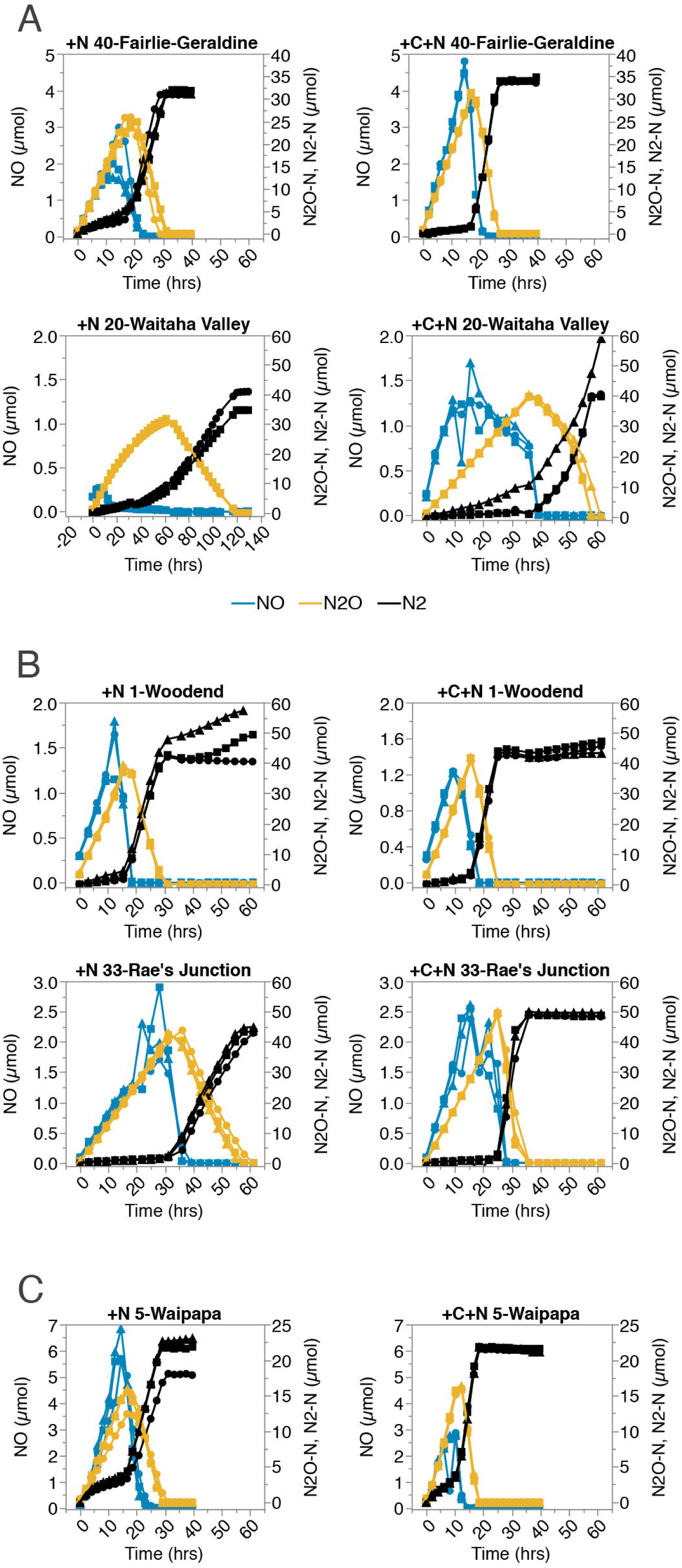
Effect of carbon (10mM Na-glutamate + 4mM NH_4_NO_3_ by flooding and draining) on soil denitrification kinetics in representative soils ranging in N_2_O hypothetically emitted (%)/phenotypes: concurrent (A), sequential (B) and intermediate (C). Triplicate incubations per treatment (dots, squares, triangles) were carried out under anoxia without oxic preincubation. Carbon amended treatments right, C negative controls left. Single leaky reps excluded for 20 +N and 40 +C+N. N_2_O (Orange), N_2_ (Black) and NO (Blue) are reported as µmol-N per vial. Carbon additions shift kinetics in the tested concurrent soils towards sequential N_2_O production/reduction and greater NO accumulation while no dramatic change is observed for the sequential or intermediate soils. Graphs with axes scaled to same maximum available (Figure S5)

Carbon amendment had variable impacts on denitrification phenotype, respiration and denitrification rates depending on the soil (Figure 5, Table 3). Added carbon clearly relieved some limitation as we observed increased CO_2_ production (Table 3) and reduced time to complete denitrification in all soils (Figure 5), but impacts on denitrification phenotype were not in line with our original hypothesis. Carbon amendments drove concurrent soils (40-Fairlie-Geraldine, 20-Waitaha Valley) towards a more sequential phenotype (increased N_2_O hypothetically emitted % and max NO accumulation compared with +N controls, Table 3) while sequential soils (1-Woodend, 33-Rae’s Junction) maintained their sequential phenotype (similar N_2_O hypothetically emitted % and max NO accumulation compared with +N controls, Table 3). Our C amended intermediate soil 5-Waipapa did not appear to respond in the same way as other soils. The soil accumulated less NO than the N amended control (Difference max NO, 3.26µmol) and showed a variable response in measures of N_2_O emission potential (Table 3).

## Discussion

The initial incubation experiment unexpectedly revealed a continuum of soil denitrification phenotypes based on the timing of N_2_O reduction/production. The most striking soils (Figure 1C) carried out N_2_O production and reduction steps almost entirely sequentially, accumulating most N as N_2_O in vial headspace before initiating rapid N_2_O reduction. In an open vial or pasture soil, this emission pattern is predicted to result in up to 100% emission of produced N_2_O, depending on soil physical properties (e.g. depth, water filled porosity) determining the ability of delayed N2O reduction to transform N2O before emission. In addition to our initial aim of re-assessing previously observed links between N2O/(N2O+N2), pH and microbial community composition (Samad *et al.*, 2016b, 2016a), we explored the potential causes of these contrasting N2O production/reduction phenotypes which are hypothesized to be due to a transient mechanism of action, potentially a reversible inhibition or regulatory process.

### 4.1 The role of pH

The correlation between low pH and high N_2_O/(N_2_O+N_2_) ratio is well documented and has been demonstrated in a variety of experimental systems (Bergaust *et al.*, 2010; Liu *et al.*, 2014; Samad *et al.*, 2016a). As such, we were surprised to find that pH was not correlated to measures of N_2_O emission potential in the present study. It is possible that variability of other factors influencing N_2_O emission ratios overshadowed a pH effect in this particular data. pH correlations with N_2_O/(N_2_O+N_2_) ratios are most often observed through variable pH manipulation within a single site or soil e.g. (Čuhel *et al.*, 2010; Liu *et al.*, 2010; Simek and Cooper, 2002), while the studied soils were from varying geographical locations with variable management.

Alternatively, unobserved pH changes before the anoxic incubation period could obscure a true correlation. Comparisons between measured soil NO_3_^-^ + NO_2_^-^ before oxic incubation and predicted NO_3_^-^ + NO_2_^-^ at the beginning of the anoxic period (based on final N_2_ accumulated) showed nitrification must have occurred in most soils while they were under oxic conditions (Figure 4A). Nitrification of ammonium results in the release of two H^+^ ions per molecule of ammonium oxidized (Rowell and Wild, 1985; Zhao *et al.*, 2014) and therefore could have caused significant acidification of incubated soils in the present study, making initial pH measurements irrelevant. Various lines of evidence seem to counter this hypothesis:

1. There was no correlation between nitrification activity, predicted from the difference in initial measured NO_3_^-^ + NO_2_^-^ vs. estimated at the start of the anoxic incubation and N_2_O/(N_2_O+N_2_) suggesting that pH changes due to nitrification were negligible or at least too minor to completely define soil pH trends (Spearmans’s correlation N_2_O/(N_2_O+N_2_) vs. NO_3_^-^ + NO_2_^-^ accumulation during oxic incubation µmol, ρ=0.15, p > 0.05).
2. Samad *et al.* (2016b, 2016a) observed a significant correlation between initial soil pH and N_2_O/(N_2_O+N_2_) using the same incubation methodology used here. Therefore, variable acidification was not an issue, even though high gaseous N accumulation suggested substantial nitrification occurred.
3. pH was still not correlated to N_2_O/(N_2_O+N_2_) in repeated soil incubations without an oxic period (and presumably minimal nitrification) (Spearmans’s correlation, ρ=0.10, p > 0.05).
4. Omission of the oxic incubation period usually resulted in increased hypothetically emitted N_2_O (%) compared with initial incubations (Table 3). The opposite would be expected if significant amounts of acidification occurred during oxic periods.

Based on these arguments, we tentatively conclude that factors other than pH were the most important drivers of N_2_O production/reduction phenotypes and N_2_O/(N_2_O+N_2_) in the current study but do not doubt that soil pH could exert effects on the observed phenotype, associated N_2_O/(N_2_O+N_2_) ratios and NO accumulation patterns based on retrospective analysis of (Samad *et al.*, 2016b, 2016a). Previous evidence suggests pH based control of N_2_O/(N_2_O+N_2_) is due to a post-trasnscriptional impairment of enzyme maturation in the periplasm (Bergaust *et al.*, 2010; Liu *et al.*, 2014), however, we suggest this does not explain well the delayed nature of N_2_O reduction observed in (Samad *et al.*, 2016a) or elsewhere (Liu *et al.*, 2010).

### 4.2 The role of microbial community composition and distal regulators in determining observed phenotypes

Samad *et al.* (2016b, 2016a) previously linked N_2_O/(N_2_O+N_2_), pH, and 16S microbial community composition using the same methodology used here. However, it remained unclear whether correlations between N_2_O/(N_2_O+N_2_) and 16S microbial community composition indicated a true causal link. Based on the well-known impacts of pH on both N_2_O/(N_2_O+N_2_) (Simek and Cooper, 2002) and microbial community structuring (Lauber *et al.*, 2009; Kaminsky *et al.*, 2017; Samad *et al.*, 2016b), a plausible explanation was that pH independently determined N_2_O/(N_2_O+N_2_) and microbial community composition. A similar 3-way correlation emerged here but with average daily rainfall at the sample sites in place of pH. Again, it is plausible that long term rainfall patterns or a linked variable separately influenced N_2_O/(N_2_O+N_2_) and community composition, however, taken together Samad *et al.* (2016a, 2016b) and the present study may indicate an alternate story: a consistent link (though less strong here than Samad *et al.* (2016a)) between N_2_O emission potential and community composition and a consistent continuum of N_2_O production/reduction phenotypes (though less obvious in (Samad *et al.*, 2016a)) both occurring across different soil sets among alternate potential confounding drivers (rainfall patterns here, pH in (Samad *et al.*, 2016b). Thus, there is some increased support for a true link between 16S community composition and N_2_O emission potential. Soil phenotypes were clearly sensitive to manipulations i.e. carbon addition (Figure 5 Left side vs. right side) and methodology changes altered N_2_O emission potential/phenotypes (Figure 1 vs. Figure 5), but different communities could hypothetically display a greater propensity for more sequential or concurrent denitrification under consistent proximal regulators. This could be due to, for example, alternate denitrification regulatory phenotypes (e.g. early *nosZ* expression) of community members between soils (Liu *et al.*, 2013; Lycus *et al.*, 2017; Bergaust *et al.*, 2011).

It remains unclear how exactly rainfall patterns shape the denitrifying community and N_2_O emission potential, though past hydrological experience has previously been linked to the timing of soil N_2_O reduction (Zhu *et al.*, 2013). Short term rainfalls prior to sampling, soil storage moisture and soil moisture content at the time of experimentation were irrelevant to N_2_O emission potential, therefore a longterm effect on chemistry or community selection is implied. Selection could involve recruitment of successful organisms from the available biosphere and, over longer periods, evolutionary adaption e.g. (Lynch and Neufeld, 2015; Parkin *et al.*, 1985). We hypothesize long periods of soil saturation, ensuing anoxia and slowed diffusion of N_2_O provide a niche in which complete and concurrent denitrifiers are more successful. Under more transient and less complete anoxia (low rainfall), denitrifiers showing short term prioritization of NO_3_^-^ + NO_2_^-^ may be selected due to a greater energy yield of NO_3_^-^ + NO_2_^-^ (Giles *et al.*, 2012; Simon and Klotz, 2013), their immediacy in the denitrification pathway, a higher availability of nitrate from nitrification coupled denitrification (due to semi-oxic conditions) (Wrage *et al.*, 2001), or the relatively poor (in comparison to prior reductases) activity of N_2_O reductase in the presence of O_2_ (Morley *et al.*, 2008).

The abundance and diversity of Clade II nitrous oxide reductase genes is previously predicted to control the N_2_O sink capacity of soils (Jones *et al.*, 2014) as nosZII carrying denitrifiers are more likely to lack N_2_O producing steps (Graf *et al.*, 2014). Although *nosZII* gene abundances and *nosZI/nosZII* gene abundance ratios here were related to pH differences (Spearman correlation N_2_O/(N_2_O+N_2_) vs. *nosZI/nosZII* ρ=-0.47, p < 0.05, vs. n*osZII* copy numbers ρ=-0.60, p < 0.05), as in previous studies (Jones *et al.*, 2014; Samad *et al.*, 2016b), they did not show any correlation to N_2_O/(N_2_O+N_2_) product ratios (Spearman’s correlation, p > 0.05) in this study, suggesting that they probably did not determine N_2_O sink capacity. Indeed, the decoupling of N_2_O/(N_2_O+N_2_) product ratios and *nosZII* abundances/ratios in this study under circumstances where pH was not found to drive N_2_O/(N_2_O+N_2_) product ratios may weaken prior claims (Jones *et al.*, 2014; Samad *et al.*, 2016b) that *nosZII* abundances affected N_2_O/(N_2_O+N_2_) product ratios rather than simply varying with the shared driver of pH.

Higher N_2_O emission potential for sequential soils occurred due to poor timing of N_2_O reduction rather than a deficit in actual N_2_O reduction capability (Figure 1), therefore a genotype based explanation (lack of *nosZ* containing denitrifiers vs. high presence of non-denitrifying nitrous oxide reducers only containing *nosZ*) for the differing N_2_O production/reduction phenotypes seems unlikely. Some of the lowest nosZII gene copy numbers were actually seen in concurrent soils with lower N_2_O emission potential (Figure 2C, e.g. 39-Lake Heron, 40-Fairlie-Geraldine, 27-Lumsden). Quantification of *nosZ* transcripts may have been more informative in the current study as it remains unclear whether *nosZ* expression was just delayed in sequential N_2_O producing/reducing soils or early function was impaired by post-transcriptional effects, which are previously observed to occlude transcription based effects (e.g. Liu *et al.*, 2010, 2014).

### 4.3 The role of proximal regulators in determining observed phenotypes

#### 4.3.1 The effect of carbon availability

Enhanced N_2_O accumulation in response to carbon limitation has been attributed to competition for electrons between the different denitrification enzymes (Pan *et al.*, 2013; Ribera-Guardia *et al.*, 2014; Dendooven *et al.*, 1994). Here, carbon additions were made to denitrifying soil incubations to test the hypothesis that sequential phenotype soils have limited electron supply and thus direct electrons preferentially towards the earlier steps of denitrification. This mechanism would also explain why impaired N_2_O reduction activity was transient i.e. as prior electron acceptors deplete, competition would be relieved. Under these circumstances, addition of carbon should lead to increased electron availability (as long as regeneration of the electron carrier pool was not already maximal) and presumably increased early N_2_O reduction. Experimental evidence here mostly contradicted that hypothesis. Carbon addition to the hypothesized “electron limited” sequential soils did not result in a consistent shift towards a concurrent phenotype, though increases in denitrification process rates do suggest that those soils were indeed somewhat carbon limited (Table 3). The overall trend observed was that carbon availability, substrate type/quality, C/N ratios or some other related effect sustained or drove soils toward a sequential phenotype with increased N_2_O hypothetically emitted % and NO accumulation, excepting soil 20-Waitaha (Figure 5). A possible explanation is that carbon addition preferentially stimulated NO_3_^-^ + NO_2_^-^ reduction leading to accumulation of NO which may in turn have an inhibitory impact on N_2_O reductase (see 4.3.2). Though it remains unclear why the initial denitrification steps would be preferentially enhanced.

Comparisons to initial soil incubations may also be informative about the role of carbon, though differences in methodology and initial NO_3_^-^ concentrations should be taken into consideration. These initial soil incubations probably had less carbon available for denitrification due carbon consumption during oxic pre-incubations as evidenced by lower CO_2_ production rates during denitrification (Table 3). If the crude assumption is made that average CO_2_ production during denitrification was proportional to carbon availability then hypothetical N_2_O emission potential in many of these soils (40-Fairlie-Geraldine, 20-Waitaha Valley, 1-Woodend) appear to exhibit a positive correlation to carbon availability (Table 3).

Based on the above observations it seems plausible that differences in carbon accounted for some of the phenotypic variation observed between soils in the original incubations. Direct measurement of starting carbon concentrations (e.g. total C, dissolved organic C) or substrates by mass spectrometry in soils would be beneficial in future investigations of the observed denitrification phenotypes.

#### 4.3.2 Nitric oxide accumulation, nitrite accumulation and nitrate concentration

Accumulation of prior N oxyanions/oxides (NO_3_^-^, NO_2_^-^, NO) can impair N_2_O reduction activity (Blackmer and Bremner, 1978; Firestone *et al.*, 1979; Gaskell *et al.*, 1981; Senbayram *et al.*, 2012; Ha *et al.*, 2015; Pan *et al.*, 2013; Zhou *et al.*, 2008; Frunzke and Zumft, 1986) due to N reductase competition for electrons e.g. (Pan *et al.*, 2013; Dendooven *et al.*, 1994) or alternate mechanisms such as direct inhibitory interaction between NO and N_2_O reductase (Frunzke and Zumft, 1986), NO_2_^-^ protonation to inhibitory nitrous acid (Zhou *et al.*, 2008) or NO_2_^-^ based enhancement of obligate N_2_O endproduct producing fungi (Maeda *et al.*, 2015). Again, transient accumulation of N oxyanions/oxides is in line with transient impairment of N_2_O reductase in sequential N_2_O producing/reducing soils. The NO accumulation patterns and timing are particularly conspicuous: NO accumulation was higher in sequential N_2_O producing/reducing soils (Spearman’s correlation, average µmol NO vial^-1^ vs. N_2_O hypothetically emitted %, ρ=0.80, p<0.0001), increased N_2_O reduction coincided with rapid depletion of NO (Figure 1C), concurrent N_2_O producing/reducing soils eventually stabilized NO levels to a low steady state (Figure 1A), and C amendments increasing or decreasing N_2_O hypothetically emitted % also increased/decreased max NO accumulation (Table 3). Based on these observations we hypothesize that sequential type soils were unable to maintain NO concentrations below an inhibitory level, resulting in impaired N_2_O reduction until NO production ceased. Alternatively, NO accumulation may be indicative of a significant NO_2_^-^ pool stimulating NO production by both abiotic and biotic processes (Lim *et al.*, 2018). NO_2_^-^ reductase is proposed to be particularly competitive with N_2_O reductase for electrons due to a shared use of the electron carrier cytochrome C550 (Richardson *et al.*, 2009; Pan *et al.*, 2013), therefore, tracking of endogenous NO_2_^-^ and evaluating responses to exogenous NO_2_^-^ in future experimentation is highly desirable.

Correlations between some measures of N_2_O emission potential and predicted NO_3_^-^ + NO_2_^-^ at the start of anoxia (Table 2) suggest initial N supply may have impacted the observed gas kinetic patterns. However, we are skeptical based on a lack of correlation with the most pertinent variables (N_2_O hypothetically emitted % and N_2_O/(N_2_O+N_2_) (max N_2_O)) and potential biases in the other measures. For instance, measures taken at the 50hr anoxia timepoint will capture higher ratios in soils with high NO_3_^-^ + NO_2_^-^ because they typically take longer to denitrify. Further, soils showed diverse relative N_2_O reduction rates at the same level of remaining NO_3_^-^ + NO_2_^-^ (Figure 4B, C).

NO_3_^-^ + NO_2_^-^ concentration effects could hypothetically be occluded in correlations between separate soils if individual soils had dramatically differing sensitivities to the similar NO_3_^-^ or NO_2_^-^ concentrations. Differing sensitivity of N_2_O reduction to NO_3_^-^ + NO_2_^-^ concentration in different soils has been previously reported, with higher sensitivity in lower pH soils (Blackmer and Bremner, 1978; Firestone *et al.*, 1979; Gaskell *et al.*, 1981). However, we did not observe pH based ranking of soils once NO_3_^-^ + NO_2_^-^ availability was accounted for (Figure S4), and this analysis cannot be considered conclusive due to the potential bias of denitrification progress; because of differences in initial NO_3_^-^+NO_2_ ^-^ different soils reached the same remaining NO_3_^-^ + NO_2_^-^ at different times and different amounts of NO_3_^-^ + NO_2_^-^ were already utilized. Experiments applying varying concentrations of NO_3_^-^ or NO_2_^-^ to pH manipulated soils or using temporally constant NO_3_^-^ concentrations (chemostats) would be necessary to understand the true impact of NO_3_^-^ and NO_2_^-^ on the observed denitrification phenotypes and N_2_O emission potential.

### 4.4 Conclusion

Here, we demonstrate considerable variation in N_2_O emission potential for New Zealand pasture soils based on the timing and activity of N_2_O reduction and associated with the accumulation of NO gas. We show an association between N_2_O production/reduction phenotypes and microbial communities in the absence of a pH effect and in conjunction with results from Samad *et al.* (2016b, 2016a) suggest this improves the plausibility of a true link between community composition and the observed phenotypes/N_2_O emission potential. Additional correlates of N_2_O emission potential/emission phenotypes are identified at both distal (long term rainfall) and proximal levels (carbon availability) which may be linked by a common mechanism of NO accumulation and inhibition. Further research on the phenomena described here should focus on directly testing the impact of NO concentrations on the observed phenotypes, the potential accumulation of NO_2_^-^ in sequential type soils, and the potential for regulatory effects such as delayed transcription of *nosZ*.

## Acknowledgements

This work was funded by the New Zealand Government through the New Zealand Fund for Global Partnerships in Livestock Emissions Research to support the objectives of the Livestock Research Group of the Global Research Alliance on Agricultural Greenhouse Gases (Agreement number: 16084 and SOW12-GPLER-OU-SM) awarded to SEM and the University of Otago, New Zealand. MH was funded by a University of Otago Postgraduate Scholarship. PD P received funding from the FACCE-ERA-GAS project MAGGE-pH under the grant agreement no. 696356. We would like to thank the Nitrogen group at the Norwegian University of Life Sciences NMBU for access to lab, robotic autosamplers, experimental and technical assistance. We also thank Steve Wakelin and AgResearch for providing historic physicochemical data and preliminary DNA samples for the analysed soils.

