## Supplementals for "Soil N_2_O emission potential falls along a denitrification phenotype gradient linked to differences in microbiome, rainfall and carbon availability"

### Supplementary Tables and Figures

**Table S1.** Basic descriptors for 20 sampled soils

| Site | pH (CaCl$_2$) | pH DNA extraction samples (CaCl$_2$) | pH DNA extraction samples ($H_2O$) | Storage moisture | $NO_3^-$+$NO_2^-$ concentration stored soils | Average rainfall (10 years) | Soil Group | Soil Sub-group | GPS Long | GPS Lat | Grazing |
|---|---|---|---|---|---|---|---|---|---|---|---|
| | | | | % | mmol/L porewater | mm/day | | | Decimal degrees | Decimal degrees | |
| 1 Woodend | 5.34 | 5.40 | 6.12 | 35% | 0.27 | 1.65 | Gley | Typic Orthic Gley | 172.6585126 | -43.34467374 | Horse |
| 3 Culverden | 5.84 | 6.03 | 6.52 | 35% | 9.79 | 1.80 | Brown | Acidic Orthic Brown | 172.8766498 | -42.77304764 | Dairy |
| 5 Waipapa | 5.71 | 5.91 | 6.67 | 23% | 7.51 | 1.76 | Pallic | Mottled Immature Pallic | 173.896982 | -42.1598611 | Sheep and beef |
| 6 Awatere Valley | 5.14 | 5.29 | 6.29 | 27% | 8.71 | 1.46 | Pallic | Argillic-fragic Perch-gley Pallic | 174.0071667 | -41.65830556 | Sheep and beef |
| 10 Cobb Valley | 5.15 | 5.25 | 6.04 | 41% | 12.61 | 5.69 | Recent | Weathered Fluvial Recent | 172.8037124 | -41.02840637 | Dairy |
| 11 Tapawera | 5.79 | 6.07 | 6.83 | 41% | 7.99 | 2.94 | Recent | Weathered Fluvial Recent | 172.8184177 | -41.37703678 | Goats |
| 18 Kumara | 5.71 | 5.89 | 6.67 | 145% | 15.08 | 7.39 | Podzol | Silt-mantled Perch-gley Podzols | 171.1595534 | -42.60513565 | Dairy |
| 20 Waitaha Valley | 5.16 | 5.00 | 6.11 | 42% | 2.97 | 8.78 | Recent | Mottled Fluvial Recent | 170.6967222 | -42.99430556 | Dairy |
| 22 Karangarua | 5.25 | 5.27 | 6.21 | 28% | 2.15 | 11.51 | Recent | Mottled Fluvial Recent | 169.9179048 | -43.48988905 | Sheep |
| 24 Makarora | 5.96 | 6.21 | 6.77 | 58% | 0.67 | 6.54 | Recent | Typic Fluvial Recent | 169.2133361 | -44.25443279 | Sheep and beef |
| 25 Crown Range | 5.39 | 5.56 | 6.35 | 45% | 5.50 | 2.10 | Recent | Weathered Fluvial Recent | 168.8868558 | -44.97856232 | Sheep |
| 27 Lumsden | 5.21 | 5.23 | 6.12 | 38% | 7.43 | 2.42 | Brown | Typic Orthic Brown | 168.4382183 | -45.72624107 | Sheep and beef |
| 30 Dacre | 5.27 | 5.72 | 6.53 | 57% | 15.52 | 3.13 | Brown | Typic Firm Brown | 168.5855143 | -46.31253048 | Dairy |
| 31 Clinton | 5.30 | 5.35 | 6.23 | 55% | 1.31 | 2.26 | Brown | Typic Firm Brown | 169.3635871 | -46.20724295 | Sheep and beef |
| 32 Glenore | 5.01 | 5.14 | 5.89 | 43% | 9.39 | 2.49 | Pallic | Mottled Fragic Pallic | 169.8841111 | -46.10494444 | Beef |
| 33 Rae's Junction | 5.06 | 5.16 | 6.01 | 40% | 5.52 | 1.66 | Recent | Weathered Fluvial Recent | 169.4684603 | -45.73937991 | Sheep and beef |
| 37 Hakataramea valley | 5.22 | 5.06 | 5.64 | 29% | 12.77 | 1.75 | Recent | Weathered Orthic Recent | 170.5911574 | -44.79071795 | Dairy |
| 38 Southbridge | 4.85 | 4.89 | 5.88 | 38% | 6.62 | 1.63 | Recent | Weathered Fluvial Recent | 172.2660278 | -43.86016667 | Sheep and beef |
| 39 Lake Heron | 4.94 | 4.67 | 5.60 | 46% | 0.20 | 3.50 | Brown | Humose Orthic Brown | 171.1658147 | -43.50402056 | Sheep |
| 40 Fairlie-Geraldine | 5.06 | 5.60 | 6.19 | 37% | 14.88 | 1.97 | Pallic | Fragic Perch-gley Pallic | 171.00175 | -44.09722222 | Sheep, beef, dairy grazing |

| Site | Total Carbon* | Total Nitrogen* | Total' Phosphorus* | C/N Ratio* | C/P Ratio* | Potential evapo transpiration * | Drainage Class |
|---|---|---|---|---|---|---|---|
| | % | % | mg/kg | | | mm | Low=poor drainage |
| 1 Woodend | 5.30 | 0.58 | 956 | 9.2 | 55.4 | 2.74 | 2 |
| 3 Culverden | 5.10 | 0.53 | 1141 | 9.6 | 44.7 | 2.89 | 5 |
| 5 Waipapa | 2.60 | 0.27 | 533 | 9.8 | 48.8 | 2.91 | 4 |
| 6 Awatere Valley | 3.80 | 0.39 | 588 | 9.8 | 64.6 | 3.10 | 3 |
| 10 Cobb Valley | 2.50 | 0.27 | 791 | 9.4 | 31.6 | 2.33 | 5 |
| 11 Tapawera | 3.20 | 0.35 | 800 | 9.2 | 40.0 | 2.37 | 5 |
| 18 Kumara | 2.60 | 0.13 | 558 | 19.9 | 46.6 | 2.12 | 2 |
| 20 Waitaha Valley | 1.70 | 0.21 | 870 | 8.0 | 19.5 | 2.18 | 3 |
| 22 Karangarua | 0.90 | 0.11 | 843 | 8.0 | 10.7 | 2.67 | 3 |
| 24 Makarora | 2.00 | 0.23 | 1181 | 8.8 | 16.9 | 2.45 | 4 |
| 25 Crown Range | 3.10 | 0.33 | 964 | 9.4 | 32.2 | 2.42 | 4 |
| 27 Lumsden | 4.50 | 0.49 | 919 | 9.3 | 49.0 | 2.18 | 5 |
| 30 Dacre | 5.70 | 0.59 | 1545 | 9.8 | 36.9 | 2.11 | 4 |
| 31 Clinton | 4.40 | 0.44 | 787 | 10.1 | 55.9 | 2.09 | 4 |
| 32 Glenore | 3.50 | 0.38 | 696 | 9.3 | 50.3 | 2.18 | 2 |
| 33 Rae's Junction | 3.10 | 0.33 | 787 | 9.3 | 39.4 | 2.34 | 5 |
| 37 Hakataramea valley | 3.80 | 0.39 | 803 | 9.9 | 47.3 | 2.03 | 4 |
| 38 Southbridge | 5.30 | 0.56 | 893 | 9.5 | 59.4 | 2.48 | 5 |
| 39 Lake Heron | 6.00 | 0.48 | 652 | 12.4 | 92.0 | 1.71 | 5 |
| 40 Fairlie-Geraldine | 3.90 | 0.40 | 793 | 9.7 | 49.2 | 1.68 | 2 |

*Data taken from 2011 samples (Wakelin *et al.*, 2013)

**Table S2.** Key variables for anoxic incubations of 20 soils amended with 2mM $NH_4NO_3$

| Site | $NO_3^-$+$NO_2^-$ concentration post flood and drain | Incubation moisture | Predicted nitrification oxic period | Predicted $NO_3^-$+$NO_2^-$ concentration post oxic period | $N_2OI$ (50hr) | $N_2OI$ ($N_2$ plateau) | $N_2O/(N_2O+N_2)$(50hrs) | $N_2O/(N_2O+N_2)$ (max $N_2O$) | $N_2O$ hypo emit | Phenotype | Max NO | Average NO | $N_2$ plateau |
|---|---|---|---|---|---|---|---|---|---|---|---|---|---|
| | mmol/L porewater | % | µmol-N/vial | mmol/L | | | | | % final N | | µmol/vial | µmol/vial | µmol N2-N/vial |
| 1 Woodend | 3.36 | 60% | 50.91 | 7.58 | 0.98 | 0.66 | 0.98 | 0.96 | 96% | Sequential | 4.25 | 1.54 | 91.50 |
| 3 Culverden | 5.37 | 53% | 88.24 | 13.64 | 0.96 | | 0.97 | 0.89 | 87% | Intermediat | 8.93 | 4.76 | 145.52 |
| 5 Waipapa | 2.96 | 42% | 22.93 | 5.68 | 0.91 | 0.64 | 0.89 | 0.84 | 87% | Intermediat | 3.25 | 1.34 | 47.91 |
| 6 Awatere Valley | 3.75 | 38% | 14.30 | 5.63 | 0.94 | 0.73 | 0.96 | 0.85 | 88% | Sequential | 2.98 | 1.35 | 42.90 |
| 10 Cobb Valley | 4.26 | 57% | 47.76 | 8.46 | 0.92 | 0.58 | 0.89 | 0.68 | 64% | Concurrent | 0.25 | 0.06 | 96.27 |
| 11 Tapawera | 3.36 | 53% | 52.14 | 8.31 | 0.92 | 0.61 | 0.90 | 0.78 | 72% | Concurrent | 2.39 | 0.49 | 87.44 |
| 18 Kumara | 11.25 | 156% | -13.43 | 10.82 | 0.84 | 0.67 | 0.79 | 0.79 | 77% | N-A | 24.26 | 5.33 | 337.07 |
| 20 Waitaha Valley | 2.58 | 50% | 2.77 | 2.86 | 0.80 | 0.57 | 0.71 | 0.71 | 74% | Concurrent | 0.21 | 0.04 | 28.48 |
| 22 Karangarua | 2.40 | 27% | 4.65 | 3.26 | 0.62 | 0.54 | 0.61 | 0.61 | 53% | Concurrent | 0.37 | 0.08 | 17.77 |
| 24 Makarora | 1.75 | 58% | 43.46 | 5.47 | 0.28 | 0.10 | 0.24 | 0.24 | 11% | Concurrent | 0.07 | 0.02 | 63.83 |
| 25 Crown Range | 3.37 | 56% | 19.18 | 5.07 | 0.84 | 0.58 | 0.80 | 0.80 | 83% | Intermediat | 1.62 | 0.64 | 57.16 |
| 27 Lumsden | 3.09 | 60% | 26.75 | 5.33 | 0.87 | 0.56 | 0.84 | 0.74 | 76% | Concurrent | 0.48 | 0.18 | 63.72 |
| 30 Dacre | 4.37 | 73% | 58.64 | 8.36 | 0.93 | 0.64 | 0.90 | 0.73 | 71% | Concurrent | 0.64 | 0.12 | 122.80 |
| 31 Clinton | 2.04 | 71% | 9.57 | 2.72 | 0.28 | 0.53 | 0.78 | 0.78 | 76% | Concurrent | 0.71 | 0.41 | 38.47 |
| 32 Glenore | 2.46 | 59% | 17.06 | 3.90 | 0.80 | 0.62 | 0.94 | 0.94 | 97% | Sequential | 0.71 | 0.50 | 46.00 |
| 33 Rae's Junction | 2.58 | 56% | 17.50 | 4.14 | 0.87 | 0.65 | 0.87 | 0.87 | 98% | Sequential | 0.92 | 0.61 | 46.38 |
| 37 Hakataramea valley | 3.03 | 52% | 28.25 | 5.76 | 0.93 | 0.76 | 0.96 | 0.96 | 109% | Sequential | 23.20 | 12.81 | 59.62 |
| 38 Southbridge | 2.84 | 57% | 24.20 | 4.95 | 0.92 | 0.59 | 0.87 | 0.73 | 80% | Concurrent | 0.66 | 0.24 | 56.66 |
| 39 Lake Heron | 1.95 | 64% | -10.63 | 1.12 | 0.26 | 0.40 | 0.56 | 0.56 | 53% | Concurrent | 0.45 | 0.28 | 14.30 |
| 40 Fairlie-Geraldine | 2.58 | 55% | 33.89 | 5.65 | 0.69 | 0.44 | 0.60 | 0.59 | 41% | Concurrent | 0.29 | 0.10 | 62.33 |

| Site | Average $CO_2$ rate (oxic) | Average $CO_2$ rate (anoxic) |
|---|---|---|
| | µmol/h | µmol/h |
| 1 Woodend | 3.45 | 1.99 |
| 3 Culverden | 2.80 | 1.08 |
| 5 Waipapa | 2.12 | 0.74 |
| 6 Awatere Va | 2.06 | 1.05 |
| 10 Cobb Vall | 3.23 | 1.18 |
| 11 Tapawera | 2.75 | 0.80 |
| 18 Kumara | 15.21 | 6.85 |
| 20 Waitaha V | 2.38 | 0.90 |
| 22 Karangaru | 1.45 | 0.52 |
| 24 Makarora | 3.05 | 0.98 |
| 25 Crown Ra | 2.80 | 1.28 |
| 27 Lumsden | 1.82 | 0.91 |
| 30 Dacre | 2.92 | 1.05 |
| 31 Clinton | 4.32 | 2.12 |
| 32 Glenore | 2.71 | 1.61 |
| 33 Rae's Junc | 2.87 | 1.34 |
| 37 Hakataram | 4.99 | 1.77 |
| 38 Southbrid | 1.79 | 0.76 |
| 39 Lake Hero | 3.31 | 1.04 |
| 40 Fairlie-Ge | 2.68 | 1.15 |

**Table S3.** Correlations between all rainfall and $N_2O$ emission metrics

| Variable | by Variable | Spearman ρ | Prob>\|ρ\| |
|---|---|---|---|
| $N_2O$ hypothetically emitted (%) | Av daily rainfall (10 years, mm) | -0.692 | 0.0007 |
| $N_2O$ hypothetically emitted (%) | Av daily rainfall (year, mm) | -0.681 | 0.0009 |
| $N_2O$ hypothetically emitted (%) | Av daily rainfall (month, mm) | -0.412 | 0.071 |
| $N_2O$ hypothetically emitted (%) | Soil moisture % storage | -0.3703 | 0.108 |
| $N_2O$ hypothetically emitted (%) | Soil moisture % incubation | -0.1370 | 0.5645 |
| $IN_2O$ (52.6hr) | Av daily rainfall (10 years, mm) | -0.568 | 0.0089 |
| $IN_2O$ (52.6hr) | Av daily rainfall (year, mm) | -0.574 | 0.0081 |
| $IN_2O$ (52.6hr) | Av daily rainfall (month, mm) | -0.390 | 0.0896 |
| $IN_2O$ (52.6hr) | Soil moisture % storage | -0.4870 | 0.0294 |
| $IN_2O$ (52.6hr) | Soil moisture % incubation | -0.2252 | 0.3399 |
| $IN_2O$ ($N_2$ plateau) | Av daily rainfall (10 years, mm) | -0.477 | 0.0388 |
| $IN_2O$ ($N_2$ plateau) | Av daily rainfall (year, mm) | -0.484 | 0.0357 |
| $IN_2O$ ($N_2$ plateau) | Av daily rainfall (month, mm) | -0.298 | 0.2149 |
| $IN_2O$ ($N_2$ plateau) | Soil moisture % storage | -0.3301 | 0.1675 |
| $IN_2O$ ($N_2$ plateau) | Soil moisture % incubation | -0.1195 | 0.6262 |
| $N_2O/(N_2O+N_2)$(50hrs max $N_2O$) | Av daily rainfall (10 years, mm) | -0.587 | 0.0066 |
| $N_2O/(N_2O+N_2)$(50hrs max $N_2O$) | Av daily rainfall (year, mm) | -0.605 | 0.0048 |
| $N_2O/(N_2O+N_2)$(50hrs max $N_2O$) | Av daily rainfall (month, mm) | -0.338 | 0.1445 |
| $N_2O/(N_2O+N_2)$(50hrs max $N_2O$) | Soil moisture % storage | -0.4509 | 0.046 |
| $N_2O/(N_2O+N_2)$(50hrs max $N_2O$) | Soil moisture % incubation | -0.1408 | 0.5537 |
| $N_2O/(N_2O+N_2)$(max $N_2O$) | Av daily rainfall (10 years, mm) | -0.608 | 0.0045 |
| $N_2O/(N_2O+N_2)$(max $N_2O$) | Av daily rainfall (year, mm) | -0.620 | 0.0036 |
| $N_2O/(N_2O+N_2)$(max $N_2O$) | Av daily rainfall (month, mm) | -0.436 | 0.0546 |
| $N_2O/(N_2O+N_2)$(max $N_2O$) | Soil moisture % storage | -0.3613 | 0.1175 |
| $N_2O/(N_2O+N_2)$(max $N_2O$) | Soil moisture % incubation | -0.1212 | 0.6106 |

**Table S4 Multiple linear regression predicting $N_2O$ hypothetically emitted (%)**

| **Summary of fit** | |
|---|---|
| RSquare | 0.314 |
| RSquare Adj | 0.185 |
| Root Mean Square Error | 0.203 |
| Mean of Response | 0.746 |
| Observations (or Sum Wgts) | 20 |

**Analysis of variance**

| **Source** | **Degrees of freedom** | **Sum of Squares** | **Mean Square** | **F Ratio** | **Prob > F** |
|---|---|---|---|---|---|
| Model | 3 | 0.300 | 0.100 | 2.439 | 0.102 |
| Error | 16 | 0.657 | 0.041 | | |
| C. Total | 19 | 0.957 | | | |

**Effect tests**

| **Source** | **# parm** | **Degrees of freedom** | **Sum of Squares** | **F Ratio** | **Prob > F** |
|---|---|---|---|---|---|
| Drainage class | 1 | 1 | 0.019 | 0.471 | 0.503 |
| PET | 1 | 1 | 0.053 | 1.295 | 0.272 |
| Av daily rainfall (10 years) | 1 | 1 | 0.235 | 5.728 | 0.0293* |

| **Expression** |
|---|
| y = 0.669 - 0.028 * Drainage class + 0.141 * pet - 0.040 * Av daily rainfall |

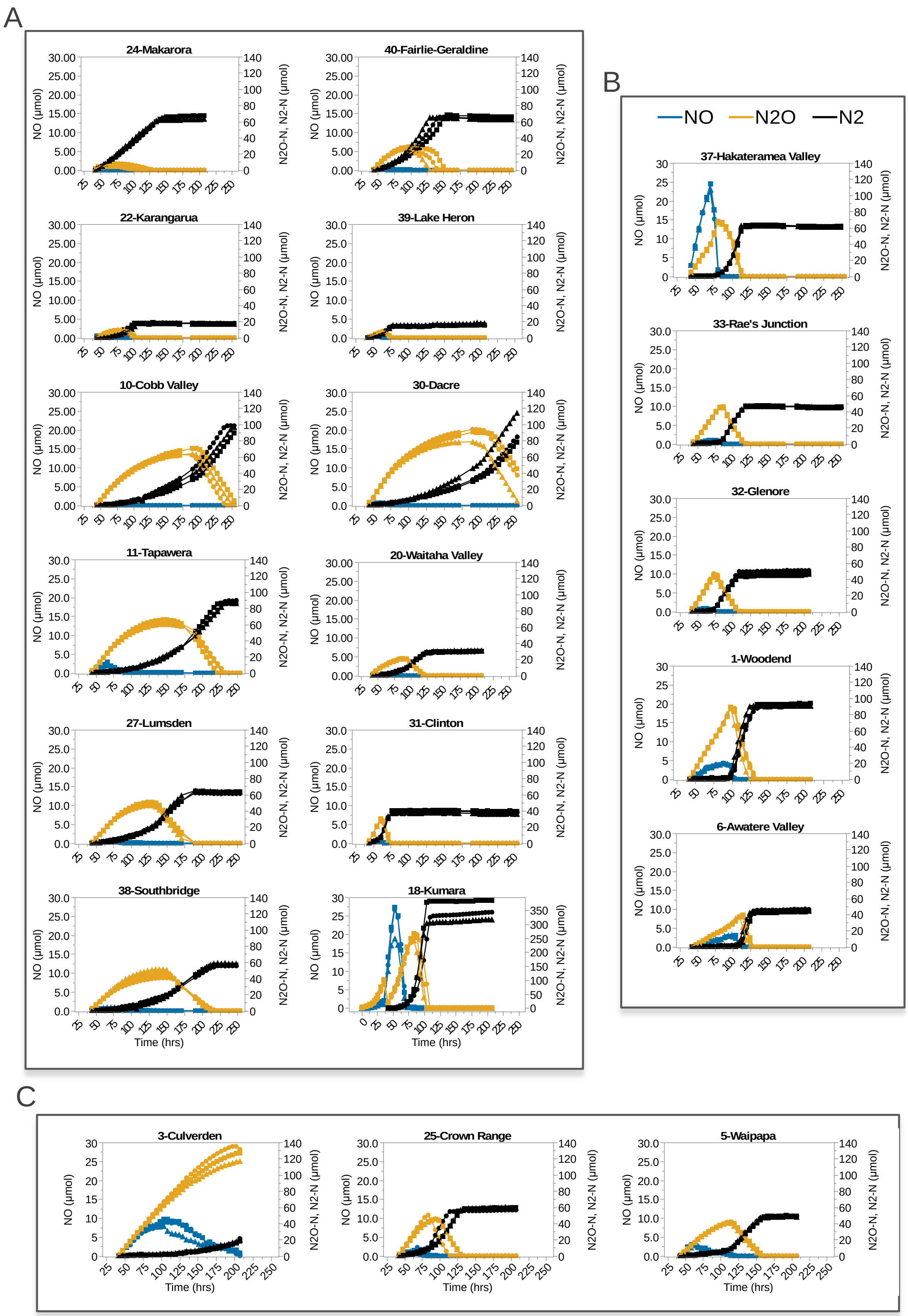
A
24-Makarora
40-Fairlie-Geraldine
22-Karangarua
39-Lake Heron
10-Cobb Valley
30-Dacre
11-Tapawera
20-Waitaha Valley
27-Lumsden
31-Clinton
38-Southbridge
18-Kumara
NO (µmol)
N2O-N, N2-N (µmol)
Time (hrs)
B
NO
N2O
N2
37-Hakateramea Valley
33-Rae's Junction
32-Glenore
1-Woodend
6-Awatere Valley
C
3-Culverden
25-Crown Range
5-Waipapa

**Figure S1.** Denitrification kinetics of all soils (as in **Figure 1**) with y-axis scaled to the same maximum. Circles, squares, triangles represent three replicate vials. $N_2O$ (Orange), $N_2$ (Black) and NO (Blue) and values are reported as μmol-N per vial.

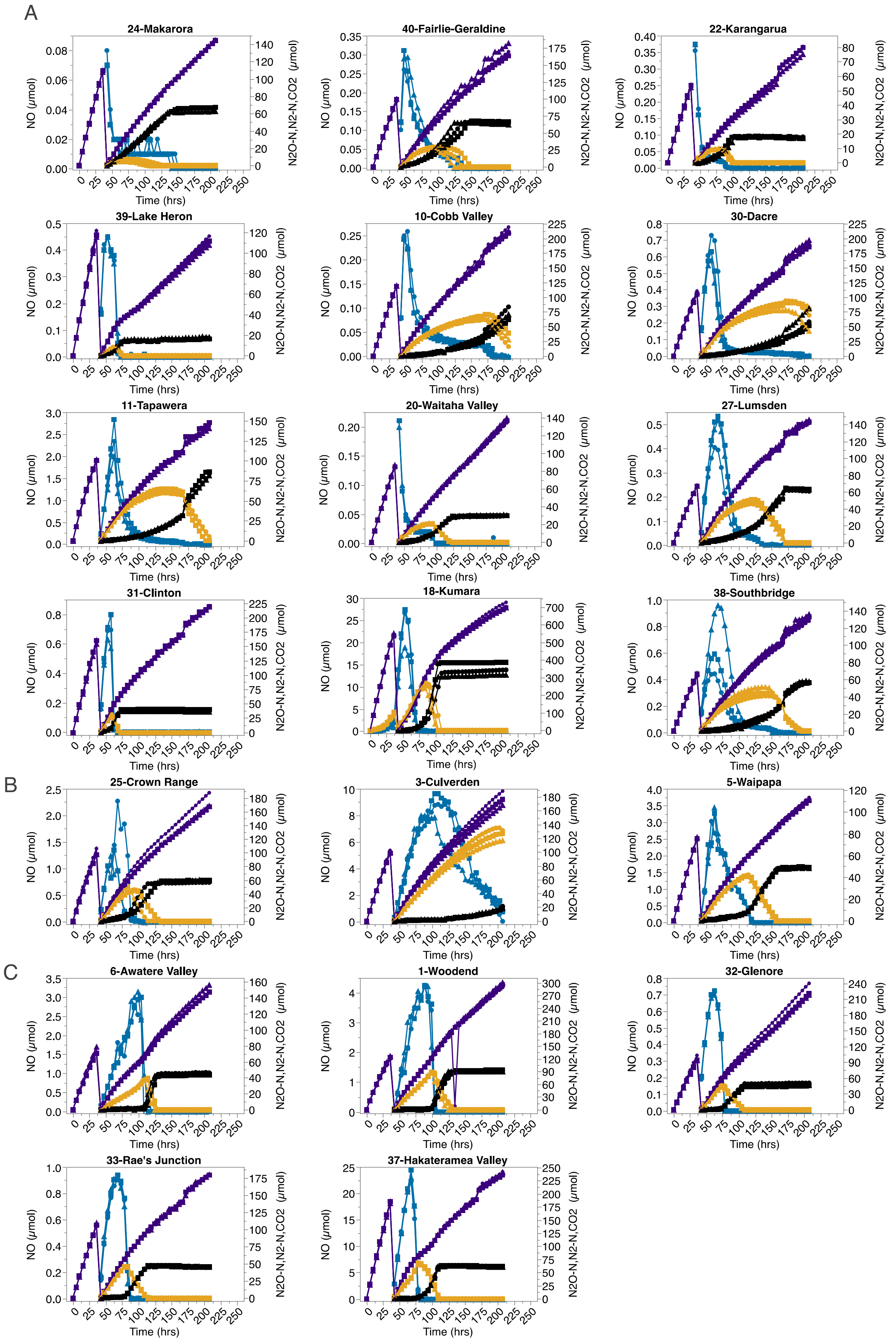

A
24-Makarora
40-Fairlie-Geraldine
22-Karangarua
39-Lake Heron
10-Cobb Valley
30-Dacre
11-Tapawera
20-Waitaha Valley
27-Lumsden
31-Clinton
18-Kumara
38-Southbridge
B
25-Crown Range
3-Culverden
5-Waipapa
C
6-Awatere Valley
1-Woodend
32-Glenore
33-Rae's Junction
37-Hakateramea Valley
NO (μmol)
N2O-N,N2-N,CO2 (μmol)
Time (hrs)

**Figure S2.** Denitrification kinetics of all soils (as in **Figure 1**) with $CO_2$ data included. Circles, squares, triangles represent three replicate vials. $N_2O$ (Orange), $N_2$ (Black) and NO (Blue) and values are reported as µmol-N per vial. $CO_2$ (purple) reported as µmol per vial.

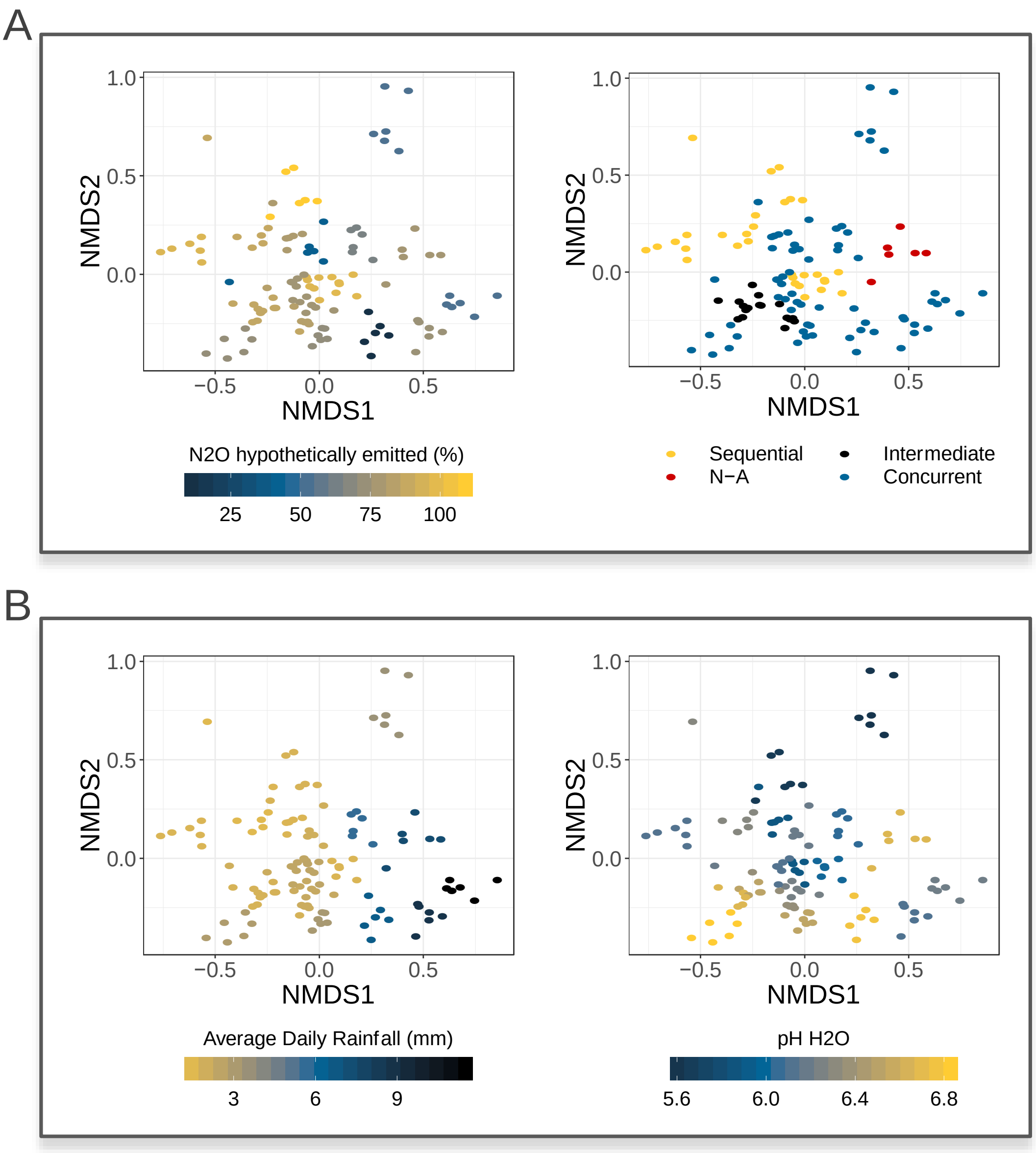

**Figure S3.** Microbial community analyses reveal links between 16S community composition and $N_2O$ emission potential/phenotypes (A), average daily rainfall over 10 years and pH (B). NMDS ordination plots (A, B) compare prokaryotic dissimilarities (Bray Curtis) of four distance specific (0m, 2.5m, 5m, 7.5m) and 2 pooled soil samples per site. Stress = 0.18. Mantel correlations of community dissimilarity vs. variables are significant for average daily rainfall over 10 years (Mantel r = 0.44, p < 0.001), pH (Mantel r = 0.41, p < 0.001) and well as $N_2O$ hypothetically emitted (%) $N_2$ (Max $N_2O$) (Mantel r = 0.26, p < 0.001).

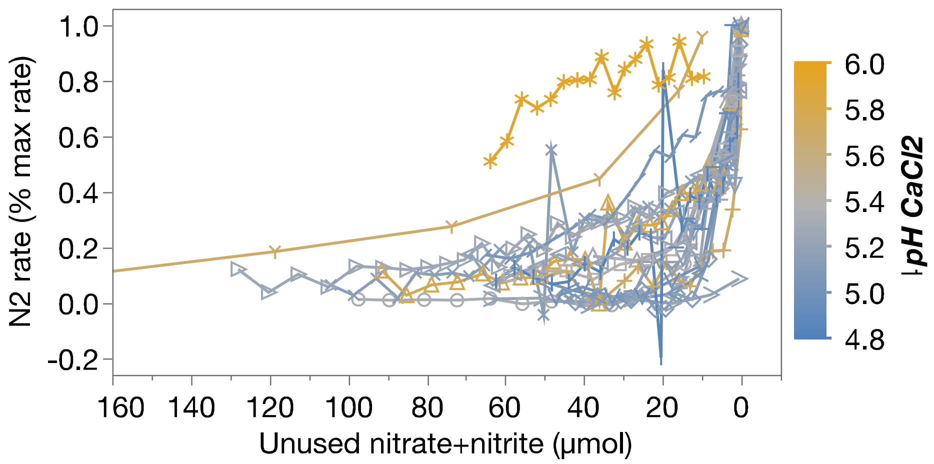

**Figure S4.** Comparison of normalized $N_2O$ reduction activity at similar $NO_3^-$ + $NO_2^-$ concentration for each soil. Ranking of soils across Y-axis could indicate potential variation in soil $N_2O$ reduction sensitivity to $NO_3^-$ + $NO_2^-$ concentration. Colour gradient shows no association between soil ranking (potential sensitivity to $NO_3^-$+$NO_2^-$) and pH.

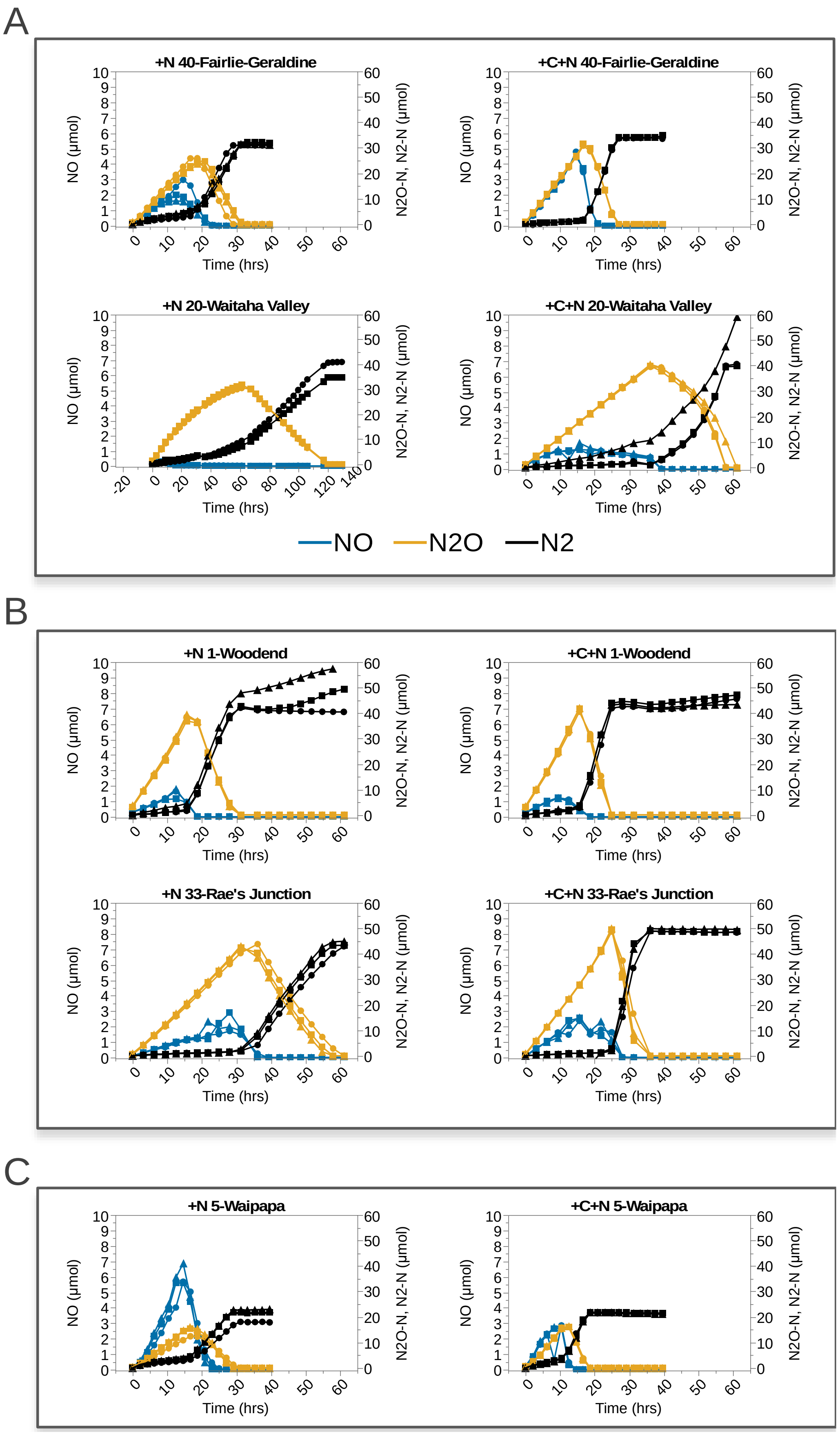

A
+N 40-Fairlie-Geraldine
+C+N 40-Fairlie-Geraldine
+N 20-Waitaha Valley
+C+N 20-Waitaha Valley
NO (µmol)
N2O-N, N2-N (µmol)
Time (hrs)
NO
N2O
N2
B
+N 1-Woodend
+C+N 1-Woodend
+N 33-Rae's Junction
+C+N 33-Rae's Junction
C
+N 5-Waipapa
+C+N 5-Waipapa

**Figure S5**. Denitrification kinetics of carbon-amended soils (as in **Figure 5**) with y-axis scaled to the same maximum. Circles, squares, triangles represent three replicate vials. $N_2O$ (Orange), $N_2$ (Black) and NO (Blue) and values are reported as µmol-N per vial.

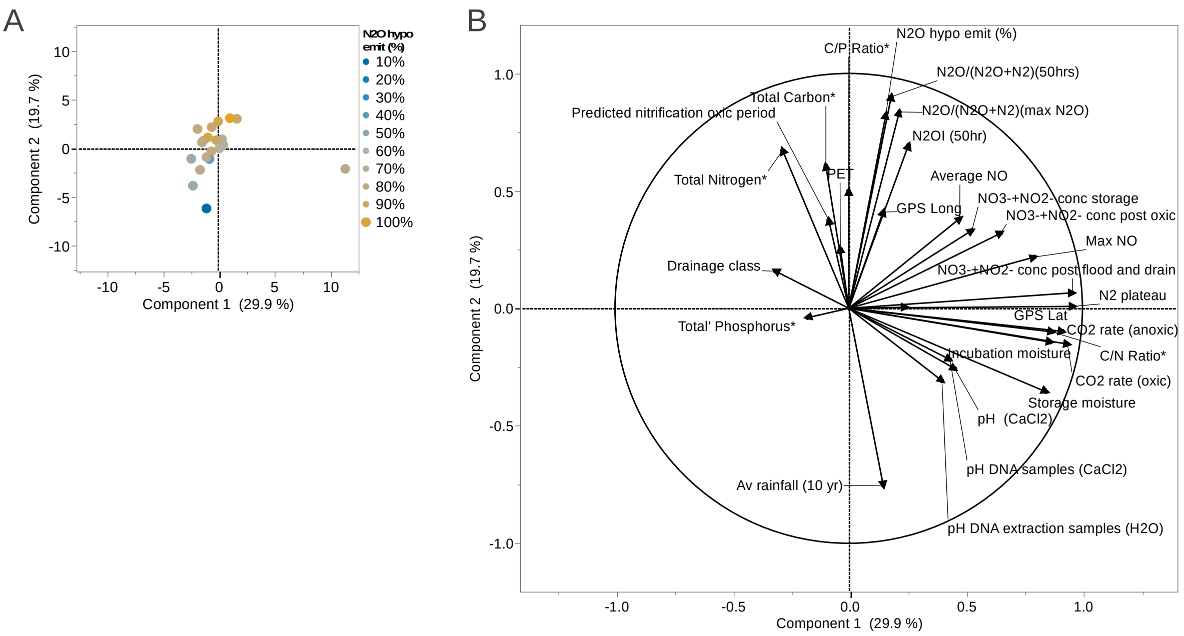

**Figure S6.** Principle component analysis of 20 soil sites (A) based on continuous soil metadata listed in Table S1 and S2. Loading plot indicating direction of factor effects (B).
